# Trans-channel fluorescence learning improves high-content screening for Alzheimer’s disease therapeutics

**DOI:** 10.1101/2021.01.08.425973

**Authors:** Daniel R. Wong, Jay Conrad, Noah Johnson, Jacob Ayers, Annelies Laeremans, Joanne C. Lee, Jisoo Lee, Stanley B. Prusiner, Sourav Bandyopadhyay, Atul J. Butte, Nick Paras, Michael J. Keiser

**Affiliations:** Institute for Neurodegenerative Diseases, University of California, San Francisco, CA, 94158, USA; Bakar Computational Health Sciences Institute, University of California, San Francisco, CA, 94158, USA; Department of Pharmaceutical Chemistry, University of California, San Francisco, San Francisco, CA, 94158, USA; Department of Bioengineering and Therapeutic Sciences, University of California, San Francisco, San Francisco, CA, 94158, USA; Kavli Institute for Fundamental Neuroscience, University of California, San Francisco, San Francisco, CA, 94158, USA; Department of Pediatrics, University of California, San Francisco, CA, 94158, USA; Center for Data-Driven Insights and Innovation, University of California, Office of the President, Oakland, CA, 94607, USA; Helen Diller Family Comprehensive Cancer Center, University of California, San Francisco, San Francisco, CA, 94158, USA

## Abstract

In microscopy-based drug screens, fluorescent markers carry critical information on how compounds affect different biological processes. However, practical considerations may hinder the use of certain fluorescent markers. Here, we present a deep learning method for overcoming this limitation. We accurately generated predicted fluorescent signals from other related markers and validated this new machine learning (ML) method on two biologically distinct datasets. We used the ML method to improve the selection of biologically active compounds for Alzheimer’s disease (AD) from high-content high-throughput screening (HCS). The ML method identified novel compounds that effectively blocked tau aggregation, which would have been missed by traditional screening approaches unguided by ML. The method improved triaging efficiency of compound rankings over conventional rankings by raw image channels. We reproduced this ML pipeline on a biologically independent cancer-based dataset, demonstrating its generalizability. The approach is disease-agnostic and applicable across diverse fluorescence microscopy datasets.

## Introduction

HCS efforts generate a wealth of complex phenotypic information that is pivotal to the drug discovery process. Extracting biological insight from these observations, however, often requires performing multiple experiments, which can demand extensive time and resources^1^. We posit that ML can extract actionable information that is otherwise encoded within HCS and learn the phenotypic relationships between related biological processes. As a corollary, we can potentially glean information on related but different biological processes—increasing the power of computationally mining large archival HCS datasets to gain new information.

The traditional way to gather more information within HCS is simply to add biological markers to track different processes. However, we cannot introduce new markers for archival datasets because the experiments are finished. Furthermore, capturing additional markers may be impractical due to expensive or cumbersome visualization procedures, like the optimization of multi-channel fluorescent immunohistochemistry or interference across detection wavelengths^2,3^. We overcame these marker limitations computationally by directly *learning* the phenotypic relationships between a highly-informative but cumbersome marker and other similar yet more easily-accessible markers. These hidden relationships were then projected into *de novo* images that displayed the desired fluorescent signal of the cumbersome marker. In this study, we focused on an archival HCS dataset that tracked the phenotypic effects of small molecules for the treatment of AD.

Despite extensive drug discovery efforts, no effective treatments exist that prevent or even slow the progression of AD^4^. Of the many therapeutic targets under investigation^5^, the pathogenic^6–13^ misfolding and accumulation of tau protein into neurofibrillary tangles (NFTs) has emerged as a target mechanism^10^. Younger decedents tend to have more aggressively propagating tau prions than older decedents^12^. Furthermore, studies have linked this tau prion propagation^14,15^ and aggregation^16^ to increased tau hyperphosphorylation. Although hyperphosphorylation is not required to form ordered assemblies of tau^17,18^, NFTs in the brains of deceased AD patients are highly enriched for hyperphosphorylated tau (pTau)^16,19^. Hyperphosphorylation inhibits the binding of tau to microtubules^20,21^, thus preventing normal microtubule assembly^21,22^ and axonal transport^23^. It may also increase the propensity of tau to be recruited into NFTs^6,23,24^. These factors promote disease phenotypes including synaptic dysfunction, enhanced neuroinflammation, and neuronal cell loss^21,23,24^.

Accordingly, discovering compounds that inhibit this tau prion propagation may improve our understanding of the disease and provide avenues for novel therapeutics. Our study focused on inhibiting the propagation and aggregation of tau in cells exposed to tau prions. To screen for compounds, we developed a HCS procedure that utilizes a biosensor cell-line overexpressing a 0N4R isoform of tau fused to the yellow fluorescent protein (YFP-tau)^25^. Due to the correlation of tau oligomerization with increased hyperphosphorylation, we posited that tracking the level of pTau in response to drug treatment would provide an additional point of validation to identify molecules that inhibit prion propagation. Furthermore, we hoped that synthesizing pTau signal would eliminate spurious signal such as contamination of the YFP channel by autofluorescent cell debris. The AT8 antibody allows us to quantify these correlated hyperphosphorylation events and is traditionally used to label disease-associated paired helical filaments in human tissue samples^26^. This antibody is specific to the Ser202/Thr205 epitope^27,28^, which is one of the most hyperphosphorylated residues in AD-afflicted brains^16,29^. Furthermore, hyperphosphorylation of this region has been linked to increased prion propagation^14,15^ and aggregation^16^. Although AT8 immunoreactivity is a useful surrogate to identify a disease-relevant phenotype and recognize clinically relevant pTau^27^, it was not used in our HCS because immunostaining would preclude some advantages of live-cell imaging^30,31^. Therefore, we turned to a different solution based on ML. We computationally synthesized a representation of the AT8 channel from existing data, rather than repeating the vast HCS with this useful antibody, which would have been laborious and cost-prohibitive.

Two seminal papers^32,33^ describe a ML method for predicting fluorescence images from bright-field images. Augmented microscopy approaches like these extract latent information from images post hoc^34^ and in a goal-driven way^35,36^. We reasoned that if two fluorescent channels were sufficiently related, we could train models to extract hidden information and generate “trans-channel” images depicting AT8-pTau from related YFP-tau images. Our logic hinged on the expectation that a ML method could both identify subtle aggregate morphology and exploit the phenotypic correlations between tau hyperphosphorylation and aggregation.

We collected a three-channel dataset—4′,6-diamidino-2-phenylindole (DAPI) nuclear marker, YFP-tau, and AT8-pTau—one year after the original HCS experiment that had only used YFP-tau and DAPI. After training our generative model on this new three-channel dataset, we applied this model to the archival HCS dataset to synthesize predicted AT8-pTau channels. We then evaluated whether the computed AT8-pTau images would better guide discovery efforts and improve the screen. To our knowledge, this is the first prospective demonstration of reliably constructing new fluorescent images *in silico* from pre-existing fluorescent markers—providing a new way to improve the drug discovery process and leverage complex biological features otherwise latent within historical HCS archives. Moreover, we assessed the generalizability of this trans-channel method by applying it to an entirely unrelated biological context of a genome-wide functional screen in osteosarcoma cells.

## Results

### We Constructed a Drug Screen Dataset of Tau Prion Propagation for ML Training

We sought to improve *in vitro* screening by constructing a new image channel from abundant, common, and relatively inexpensive channels—even after a screen’s completion. The archival HCS data contained extensive information on many compounds, but it only contained YFP-tau and DAPI images and was missing the valuable but cumbersome AT8-pTau channel. Therefore, our first task was to generate a completely independent dataset to train a ML model to predict the AT8-pTau channel solely from the YFP-tau and nuclear DAPI channels.

We constructed the new training dataset using a fixed cell system and collected all three channels (YFP-tau, DAPI, and AT8-pTau) across six 384-well plates representing a range of conditions. Ideally, ML models leverage large and diverse datasets during their training. Hence, we collected varied image data so that the model could learn to capture information about different conditions and perturbation scenarios that may be found in the HCS dataset, which itself has many different compounds present. In the training dataset, we curated a range of small molecule perturbations, each resulting in different tau aggregation and cell viability phenotypes. We used six different compounds to capture three different phenotypic spaces: reducing intracellular tau aggregation without affecting cell viability; reducing tau aggregation and reducing cell viability; and increasing tau aggregation and reducing cell viability. See Methods for a detailed description of the drug perturbation scenarios.

We constructed the three-channel training dataset to model infectious tau propagation similar to the archival HCS dataset (Methods). HEK293T cells were transfected to overexpress a YFP-tau mutant containing a familial mutation (P301S) which increases the propensity of tau to form prions that aggregate^25,37^. We then seeded the cells with an infectious mouse brain lysate passaged once through HEK293T cells expressing the microtubule-binding repeat domain of human tau with the P301L and V337M mutations. We immediately treated the cells with small molecule compounds, fixed the cells, and initially introduced the monoclonal AT8 antibody to measure phosphorylation events. We imaged three channels—DAPI, YFP-tau, and AT8-pTau (Figure 1)—to obtain 57,600 non-overlapping images that were 3 × 2048 × 2048 pixels in size. The archival HCS dataset was similar, differing predominantly in its use of live-cell imaging without AT8, and also containing thousands of compounds. Otherwise, the protocols were identical (Methods).

**Figure 1:**
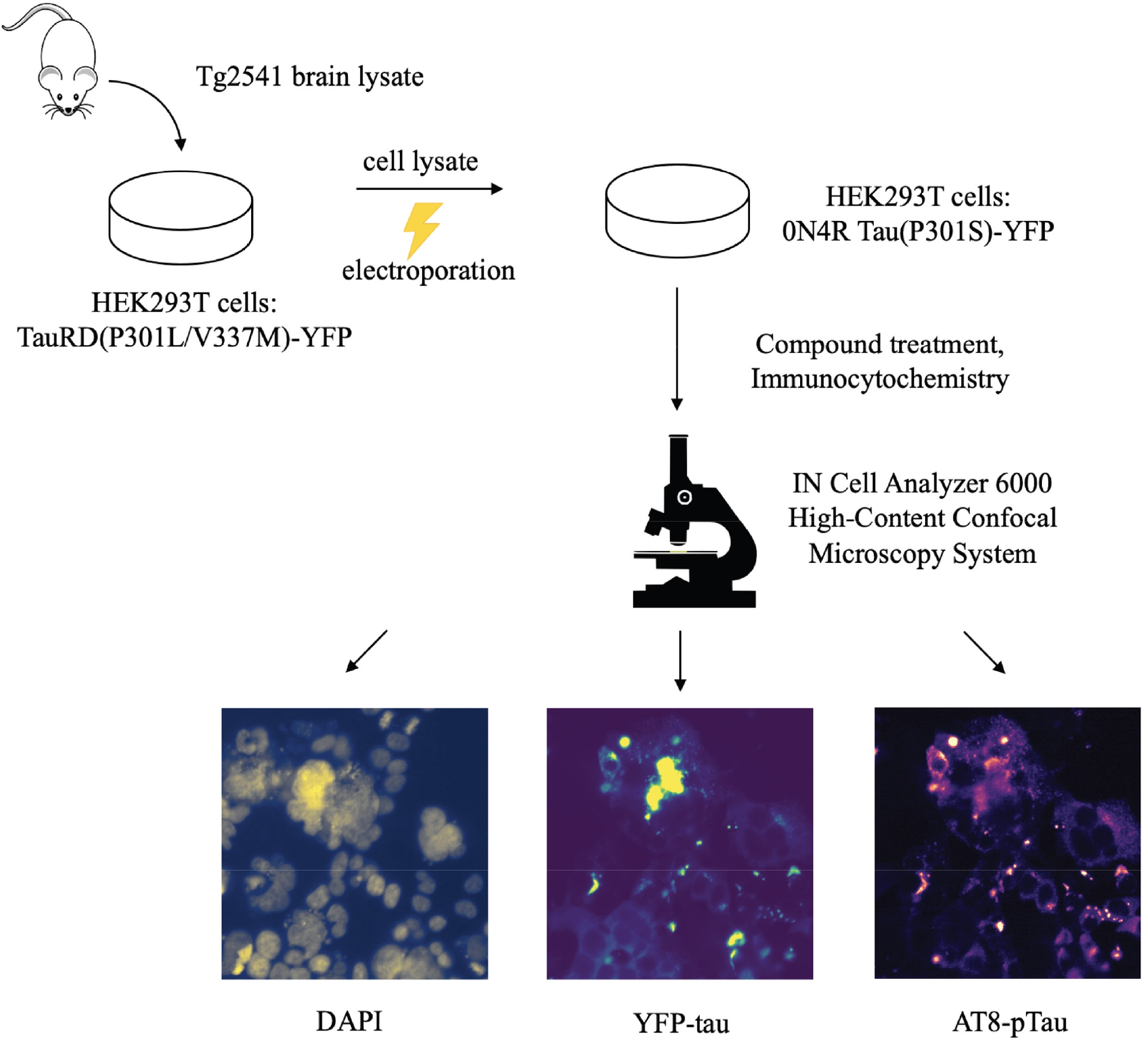
We constructed a cellular Tau(P301S)-YFP aggregation assay that modeled prion propagation *in vitro* with an additional AT8-pTau channel for ML training. An overview of the experiment used to model the adverse propagation and aggregation of tau. We used brain lysate from a tauopathy mouse model to infect a human cellular model of mutant tau that has a higher propensity to aggregate. The experiment yielded three channels used for training the ML model: DAPI, YFP-tau, and AT8-pTau. Images were enhanced with ImageJ’s auto-enhance feature solely for visualization purposes.

### We Trained a Trans-Channel ML Model to Predict Hidden Tau Phosphorylation

To learn a mapping from the input DAPI and YFP-tau channels to the output AT8-pTau channel, we developed a custom convolutional neural network (CNN) model motivated by the U-Net architecture^35^, but for a task other than segmentation. Our design employed “skip connections”^35^ that preserved lower complexity features from earlier layers and combined them with higher complexity features from deeper layers (Figure 2A; Methods). The model is 12% the size of U-Net, preserves image dimensions, and generates a non-binary image channel (Supplemental Figure 1; Methods). We made our PyTorch code and fully trained models openly available (Methods). We randomly stratified the three-channel images into a 70% train and 30% held-out test split (Methods). On visual inspection, the ML model removed aberrant signal and accurately mapped the phenotypes of the related channels to construct realistic AT8-pTau images with a high resemblance to the actual objectives (Figure 2B).

**Figure 2:**
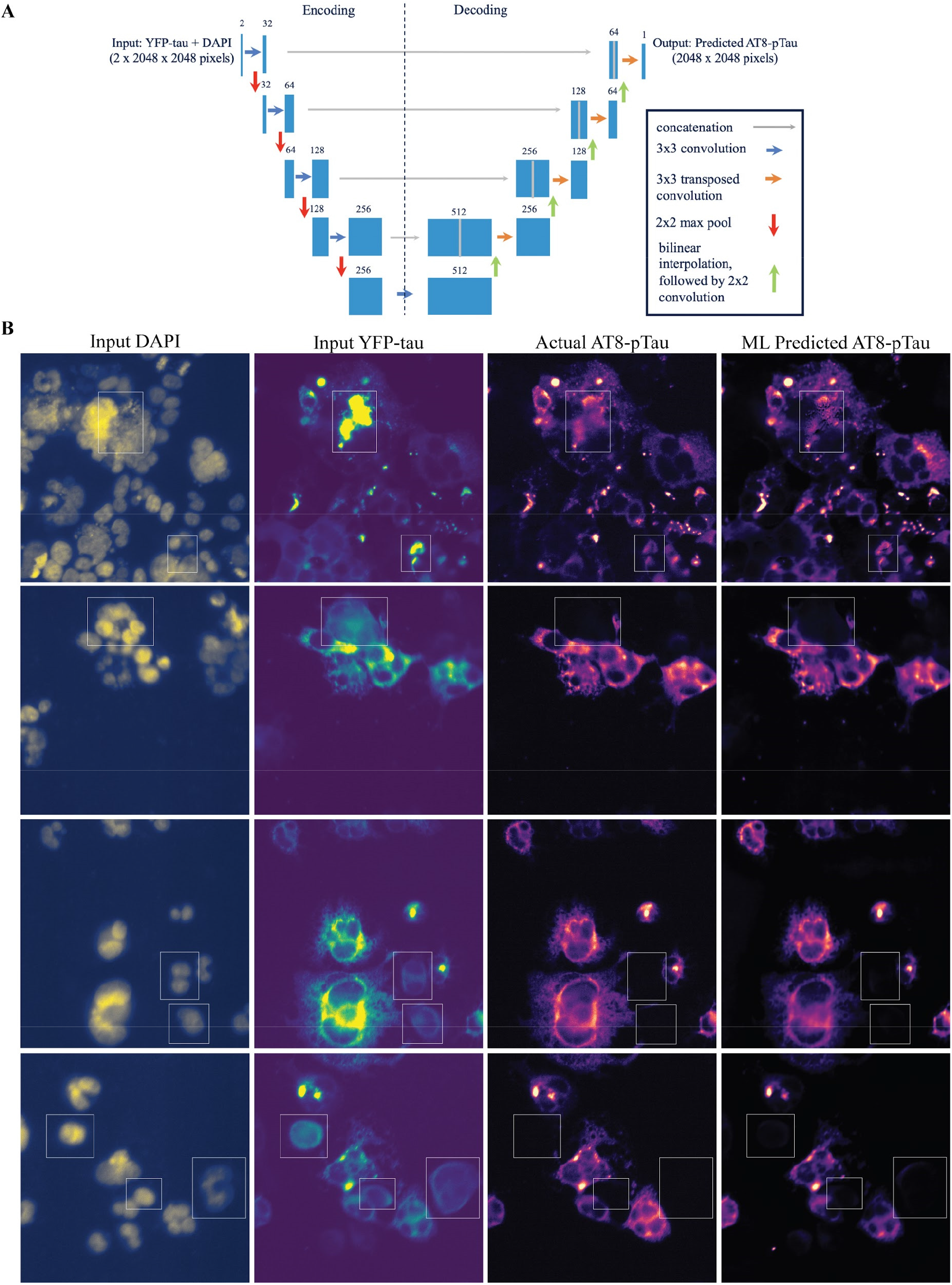
The ML model generated a phosphorylated AT8-pTau channel, solely from DAPI and YFP-tau channels. (a) Given two image channels (DAPI and YFP-tau) as input, the neural network predicted the desired AT8-pTau image. The architecture, motivated by U-Net^35^, comprised two phases: an “encoding” half which increased the cardinality of convolutional filters while reducing the image dimensions, and a “decoding” half which decreased the cardinality of filters while upsampling the image dimensions with bilinear interpolation and learnable transposed convolutions. The numbers above each box indicate the number of filters used. (b) From the input DAPI image (left column) and the input YFP-tau image (second column), the *in silico* predicted AT8-pTau image (rightmost column) is shown versus the actual, hidden AT8-pTau image (third column). White boxes mark where the AT8-pTau images most diverged from YFP-tau on visual inspection. These held-out test images were not seen during model training. For visualization purposes only, images were cropped to 512 × 512 pixels, auto-enhanced using ImageJ, and colored using the python library Matplotlib. Pearson correlation values between the predicted and actual images displayed are 0.88, 0.92, 0.93, and 0.86 from the top row to the bottom.

The ML model learned to identify and enhance regions of interest in the YFP-tau channel, namely tau phosphorylated at the Ser202/Thr205 epitope. The model learned to effectively generate the AT8-pTau images by gleaning visual information about the relationships between a hidden phosphorylation readout and the YFP and DAPI channels. It removed YFP-tau signal that was not present in the AT8-pTau image and selectively retained pertinent signal (interesting regions are boxed in Figure 2B). On visual inspection, there did not appear to be an intuitive set of visual criteria or easily predefined attributes like pixel intensity by which a human could accomplish the same task (e.g, see Supplemental Figure 2). The model’s decision-making appeared to be non-trivial.

We evaluated the model quantitatively on the held-out test set of YFP-tau and DAPI inputs and AT8-pTau objectives. To assess image similarity between the observed and predicted AT8-pTau images, we calculated the Pearson correlation and also the mean squared error (MSE) metric, which is more commonly used in regression tasks. We favored Pearson because it is invariant to pixel intensity scaling: a Pearson correlation coefficient of 1.0 indicates exactly identical images (with a possible constant scaling factor), whereas 0.0 indicates a complete pixel-wise disagreement. We obtained an average Pearson correlation of 0.74 ± 0.19 over the test set of *n*=17,280 images (Figure 3A). This corresponded to a MSE of 0.52 ± 0.39 after normalizing to zero-mean and unit-variance (Figure 3B). To quantify its pixel-wise performance, we measured how well the model balances true positive pTau pixels with false positives, assessing the error rate using the area under the curve of the receiver operating characteristic (AUROC). The model achieved an AUROC = 0.96 (Figure 3C). Importantly, we found that learning from either single channel alone was insufficient to glean the AT8 signal (Supplemental Figure 3).

**Figure 3:**
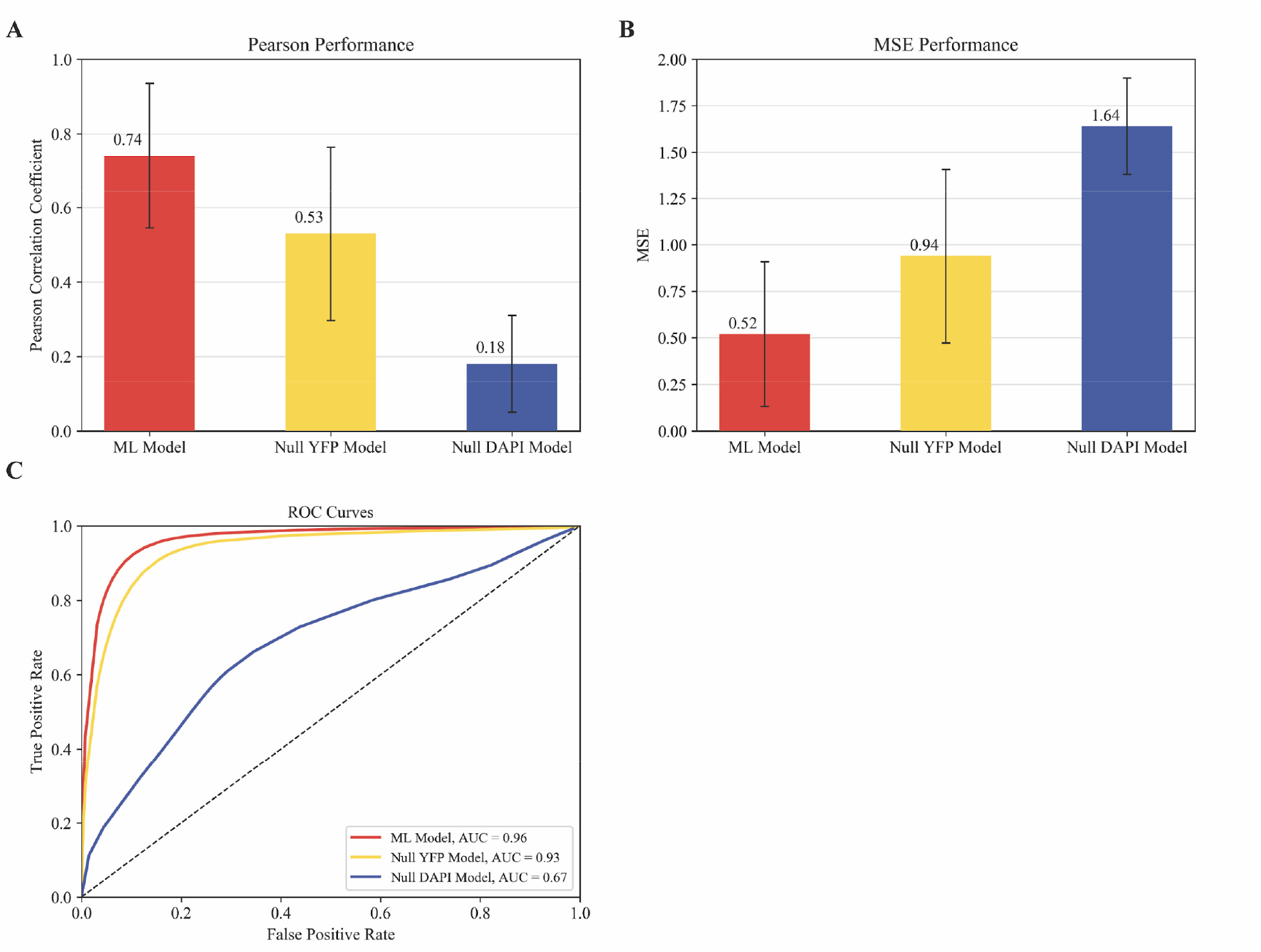
The ML model achieved good performance. (a) The average Pearson performance over the held-out test set for three models: the ML model, versus two null models derived from each of the input channels respectively. Error bars show one standard deviation in each direction centered at the mean. Assessing the ML model’s superior performance over each of the null models resulted in statistical significance of p<<0.00001. (b) Equivalent MSE results after normalizing all images to zero-mean and unit-variance (Methods). (c) The receiver operating characteristic (ROC) curves for the ML model (red) exceed the Null YFP Model (yellow) and the Null DAPI Model (blue). A binary pixel threshold of 1.0 was used to binarize the label image, while the predicted images were assessed across a range of pixel thresholds (Methods).

The YFP-tau and AT8-pTau channels contained substantial overlapping information. If they did not, the prediction task would be difficult or impossible. In theory, signal from the YFP-tau is a superset of the AT8-pTau signal and includes all of the introduced tau and aggregated protein (Supplemental Figure 4). Thus, given the similarity between the YFP-tau channel and AT8-pTau channel, we might theoretically achieve good “prediction” by simply extracting the input YFP-tau *as-is* and stipulating this as the model’s predicted output. This trivial Null YFP Model required no ML, and yielded a Pearson correlation of 0.53 ± 0.23 (corresponding to an MSE of 0.94 ± 0.47) over the entire dataset (Figure 3A-B). The ML model exceeded this baseline with an increase in average Pearson correlation of 0.21 (p<<0.00001; corresponding to a decrease in average MSE of 0.42), which is consistent with the model learning to generate an output that more closely approximated the phosphorylation state than was already provided by the YFP-tau input alone. We also created a Null DAPI Model that simply returned the input DAPI image as the output; this model obtained an average Pearson correlation of 0.18 ± 0.13 (corresponding to a MSE of 1.64 ± 0.26). The ML model’s test performance was consistent across all six drug perturbations (Supplemental Figure 5).

As with most multichannel imaging studies, we were wary of image bleed-through that the model might exploit as a hidden crutch. Excitation and emission plots of the different fluorophores are shown in Supplemental Figure 6A. To test the hypothesis that the ML model leveraged undetectable but pernicious hidden AT8 signal within the input channels, we performed pixel-intensity ablations on the input images to eliminate potential low-intensity bleed-through signal and then assessed performance (Supplemental Figure 7). We did not detect significant image bleed-through as a confounder. Accordingly, we note that any confounding signal augmenting a model’s performance would be of no help when applied subsequently to the archival HCS dataset, since the archival HCS did not undergo immunohistochemistry preparation and had neither AT8 antibody nor fluorophore.

### Trans-Channel ML Improved Hit Rate and Compound Triaging in Tauopathy HCS

Turning to its prospective use in drug discovery, we applied the trans-channel ML model to the archival HCS dataset and found practical improvements. For most HCS pipelines, immunostaining is not normally employed due to batch variability and increased labor and cost, and must be balanced against the advantages of live-cell formats such as time-course data collection. If ML could inexpensively infer these immunohistochemistry channels, then it could enhance HCS efforts by providing previously unavailable channels and inferences—thereby improving drug discovery pipelines. Similarly, one might imagine a wealth of data could be mined in a hypothesis-guided way from substantial existing datasets of completed screens in the public and private sectors, thus advancing biological knowledge and improving medical therapies.

Compounds are conventionally ranked by priority, favoring those that lower tau aggregation for secondary dose-response testing. Medicinal chemists also consider chemical structure when prioritizing. Instead, we took a naive approach by having the ML model make decisions based solely on cellular phenotype—unguided by human intuition.

To assess the ML method for practical use, we directly compared it to the conventional method of drug discovery, which was unguided by ML. To our knowledge, this study is the first practical application of trans-channel ML image generation actively used in a conventional in-house screen. Using data from the archival HCS dataset that was devoid of immunohistochemistry procedures, we predicted an AT8-pTau channel for each compound using the trained model. Due to the large dataset size, we randomly selected one run of the HCS (consisting of 1,600 unique compounds) for our analysis. We constructed a ML-derived priority queue (PQML), ranking compounds based on aggregation scores using the predicted, machine-learned AT8-pTau images. We calculated aggregation scores using cellular image software provided by General Electric (Methods), as we did in the original archival HCS evaluation pipeline a year before. The conventional method’s priority queue (PQC) ranked each compound solely by the aggregation scores of pre-existing YFP-tau images. Hence each compound received rankings in both queues and these rankings could disagree substantially.

We prospectively collected complete dose-response profiles for the top 40 compounds from each queue. We chose 40 compounds (per Methods) for each queue due to cost and labor limitations. Despite operating solely on a computationally-generated channel, the ML queue’s hit rate was comparable with the conventional method and effectively proposed overlooked compounds that passed dose-response testing. Of the 40 compounds tested from PQML, 11 passed secondary testing (27.5%). Of the 40 compounds tested from PQC, 12 were active (30%). Importantly, compounds that were highly prioritized by both the ML and conventional methods obtained a much higher hit rate than either method alone. Ten compounds ranked within the top 40 of both queues and 6 out of these 10 were confirmed by dose-response. Taken together, progressing compounds by this combined criterion yielded a success rate of 60%—an improvement of 30% over the conventional method and of 32.5% over the ML method (see Table 1). However, our sample size was smaller for these overlap compounds, which must be considered when assessing generalizability.

**Table 1:** Comparison of dose-response success rates using different methods. The Random Baseline is approximated from a previous screen, in which 26 active compounds were discovered in a screen consisting of 60,851 compounds.

| Method | Dose-Response Success Rate |
| --- | --- |
| Random Baseline | 0.043% (26 / 60,851) |
| Prospectively test top 40 compounds in PQC | 30% (12/40) |
| Prospectively test top 40 compounds in PQML | 27.5% (11/40) |
| Test compounds that are shared by both PQC's top 40 and PQML's top 40 | 60% (6/10) |

Interestingly, 5 of the 11 active compounds in PQML’s top 40 were effectively missed in the PQC, which ranked them 539^th^, 545^th^, 560^th^, 582^nd^, and 1,596^th^ out of 1,600 possible compound ranks. The activity of one such compound is shown in Figure 4A. The ML model eliminated YFP-tau aggregate signal that had falsely lowered these compounds’ priority in the PQC (Figure 4B). Supplemental Table 1 shows all compound dose-response profiles for reducing tau aggregation and cell count.

**Figure 4:**
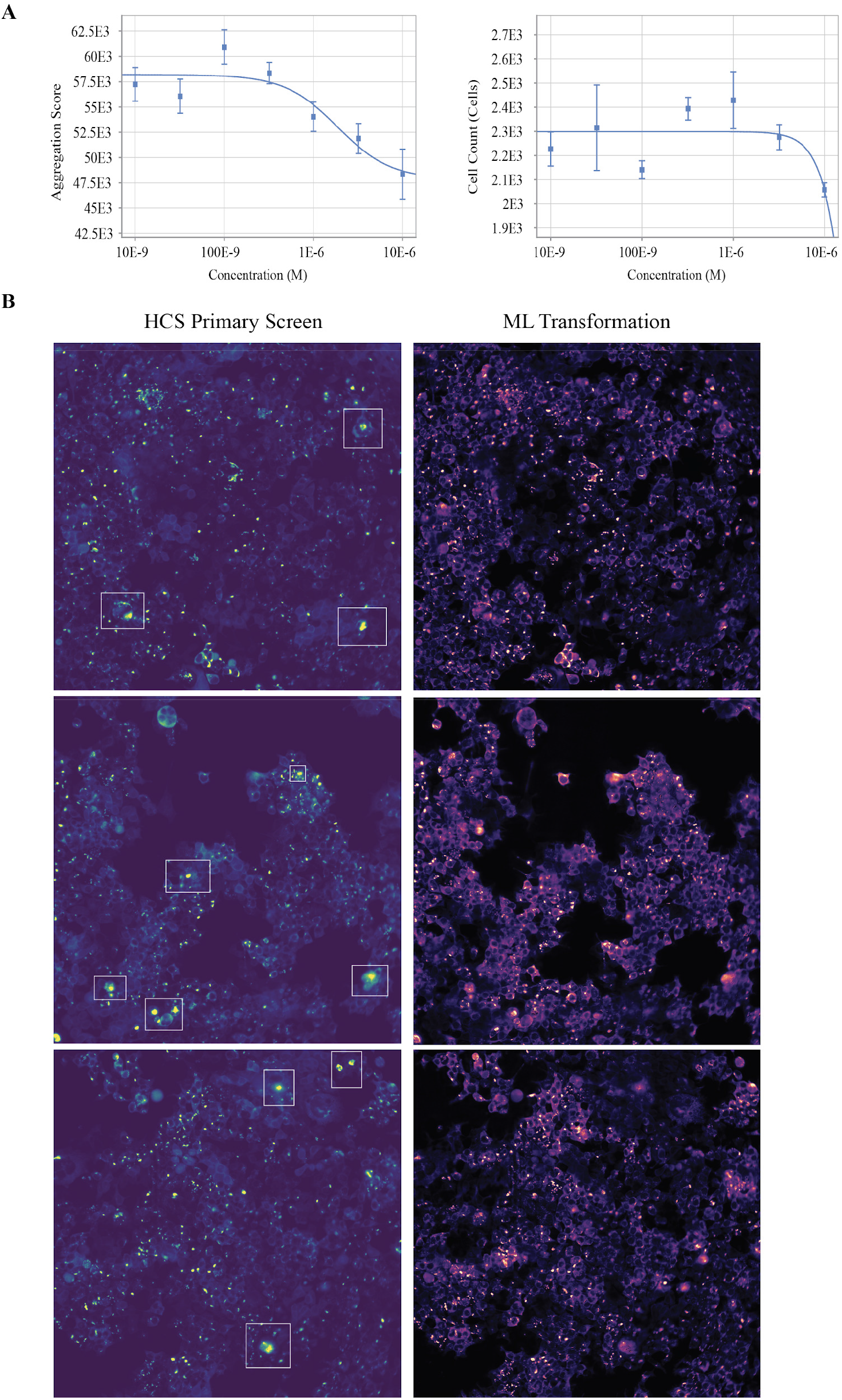
Transforming the archival HCS data with ML-predicted AT8-pTau images revealed previously unknown active compounds. (a) We rescored a plate consisting of 1,600 compounds from the archival HCS, and prospectively collected dose-responses for compounds that resulted in the lowest tau aggregation according to either PQML or PQC. Example dose-response (left) and cell count assay (right) of an active compound. Although an indicator of cell viability, cell count is not a perfect metric for toxicity, since it is possible for the compounds to inhibit the innately prolific nature of HEK293T cells. These curves are drawn from the compound DRW545, which was ranked well by the ML method (22^nd^), but poorly (545^th^) by the conventional method. (b) Example images for active compounds that the ML-based rescoring rescued. Top row: Compound DRW1596 (ranked 1596^th^ in PQC versus 14^th^ in PQML), Middle: compound DRW560 (ranked 560^th^ in PQC versus 15^th^ in PQML), Bottom: compound DRW545 (ranked 545^th^ in PQC versus 22^nd^ in PQML). YFP-tau images from the archival HCS dataset (left column) differed from the ML-predicted AT8-pTau images used for rescoring the HCS (right column), with white boxes highlighting example regions. In these regions of interest, the ML predicted the aggregate signal as not being phosphorylated at the residue of interest. We auto-enhanced and colored images with ImageJ and Matplotlib solely for visualization.

Additionally, the ML method achieved higher enrichment^38^ for active compounds. We discovered a total of 17 unique dose-response confirmed compounds from both lists. For these compounds, PQML had an area under the enrichment curve of 0.93 versus PQC’s area under the enrichment curve of 0.85 (Figure 5A). These active compounds achieved an average rank of 119 in PQML, which was better than nearly half their average rank of 235 in PQC (Figure 5B). Therefore, fewer compounds from PQML would need to be tested to find the same number of active compounds from PQC.

**Figure 5:**
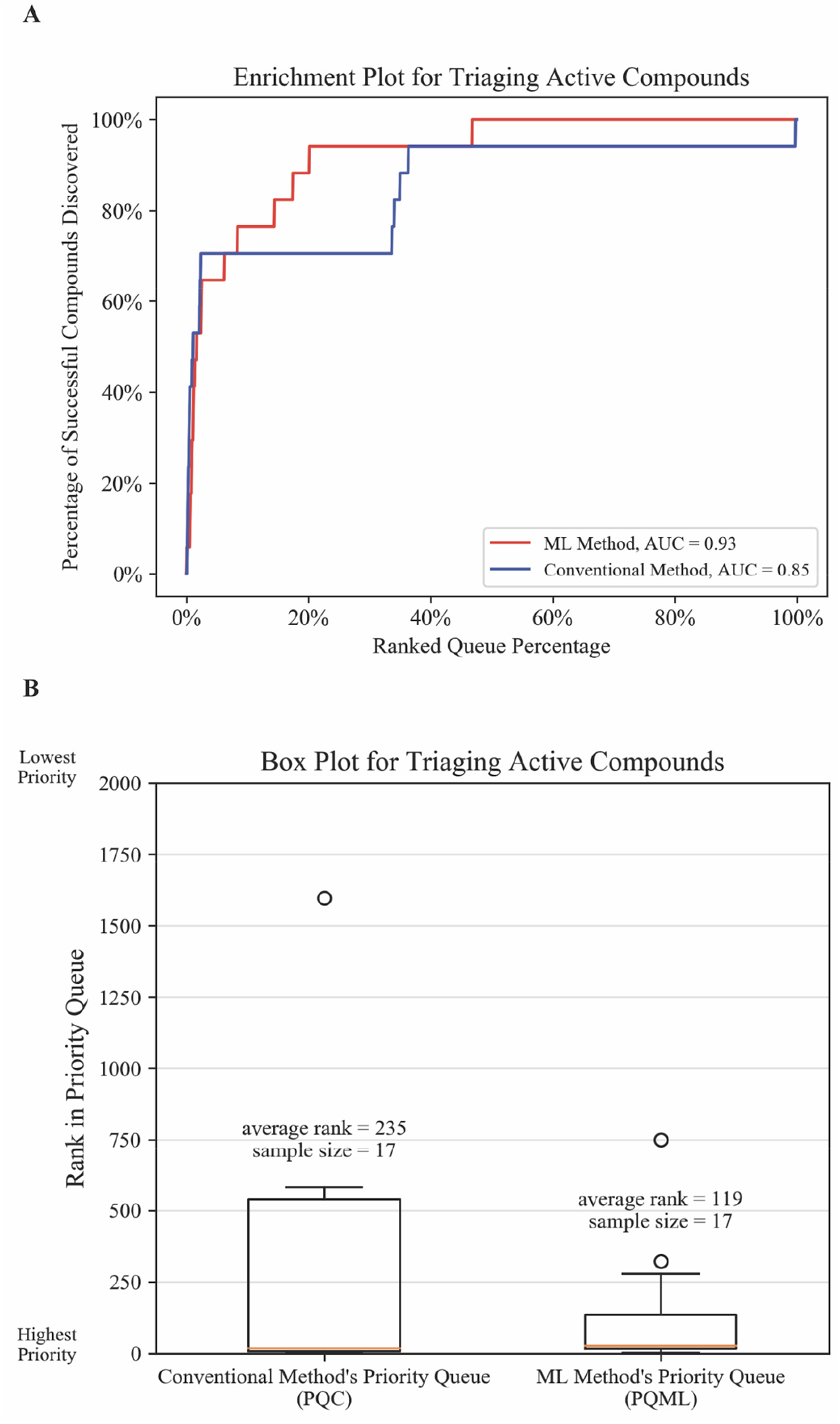
ML triaged active compounds better than the conventional method. (a) Enrichment plots comparing the conventional compound testing method with prospective, ML-guided dose-response assays. We considered the set of all active compounds (*n*=17). The y-axis shows what fraction of active compounds were discovered in the top x% of the ranked priority queue (x-axis) (100% corresponds to rank 1,600). The ML method’s larger area under the curve (AUC) indicates that its compound ranking achieves a higher success rate on prospective dose-response testing. (b) Box plots of rank accounting for all active compounds. The median line is in orange, whiskers are set at 1.5x interquartile range, and outliers are anything outside the 1.5x interquartile range (plotted as dots). PQML on average prioritizes active compounds about twice as well as PQC.

Of the 17 total active compounds from both lists, 5 fell outside of PQC’s top 40, and 6 fell outside of PQML’s top 40. However, to recover the active compounds missed in each case, we would need to test more than twice as many candidate compounds on average (*n*=764) by the PQC rankings as would be necessary by the PQML rankings (*n*=302). Hence, the ML method ranked missed-but-active compounds more than twice as well as the conventional method— including them in a smaller, more tractable search space (Supplemental Figure 8A-B). Furthermore, when we derived a compound’s dose-response activity from the ML-derived AT8-pTau images instead of scoring all queues solely by the conventional YFP-tau images, the ML method exceeded the conventional method with larger margins (Supplemental Figure 8C-F).

### We Tested Trans-Channel ML Method on an HCS with Different Biological Conditions

To assess whether the success of our trans-channel ML approach was specific merely to the tauopathy compound HCS, we investigated whether the method of learning fluorescent signal from related markers applied to a HCS unrelated in its biology and perturbagen. We performed the same trans-channel fluorescence learning task. However, compared to the prion tauopathy screen, this cancer-based HCS used an entirely different cell line, microscope, perturbation method, and fluorescent markers. ML models can exploit dataset-specific patterns in subtle ways; thus, our goal was to assess the approach’s generalizability by applying it where many of the data parameters differed from the tauopathy screen.

We performed a functional genomics HCS in a cancer cell model. In this fluorescence-based arrayed whole-genome screen (details in Methods), we plated an osteosarcoma cell line (U2OS) with a perturbation scheme that systematically knocked down all coding genes with siRNA. We marked cellular DNA with Hoechst fluorescent dye and tracked the cyclin-B1 protein with a green fluorescent protein (GFP) fusion. These two markers are biologically related, with cyclin-B1 specific to the G2/M phase of mitosis and also involved in DNA damage repair^39^. In theory, the two markers contain shared information, so we investigated whether the ML model could learn this signal relationship like it did in the tauopathy dataset. We collected a total of 324,989 images.

Unlike the tauopathy dataset, we observed subtle bleed-through signal in the raw Hoechst channel (Figure 6A; see Supplemental Figure 6B for excitation and emission spectra). Hence, as a preprocessing step to mitigate bleed-through, we ablated all pixels below the 95^th^ percentile in our Hoechst images, which inevitably removed some Hoechst signal as well (Figure 6A). We performed threefold cross-validation to predict cyclin-B1 signal solely from the ablated Hoechst channel. These models achieved an average pixel-wise Pearson correlation for channel prediction of 0.75 ± 0.13 (corresponding to an MSE of 0.50 ± 0.26) on n=108,330 test images (Figure 6B). This was 87.5% (p<<0.00001) higher than the Null Model that used the ablated input as its prediction (Pearson correlation of 0.40 ± 0.14; MSE of 1.21 ± 0.28). Despite significant ablations to the models’ input, the models were able to learn the non-trivial cyclin-B1 phenotype. Upon visual inspection, predicting where high-intensity cyclin-B1 signal resided solely from the Hoechst signal did not appear obvious. As an unexpected benefit, the ablation procedure resulted in the model predicting images that were mostly free of signal artifacts in the cyclin-B1 channel (Figure 6A).

**Figure 6:**
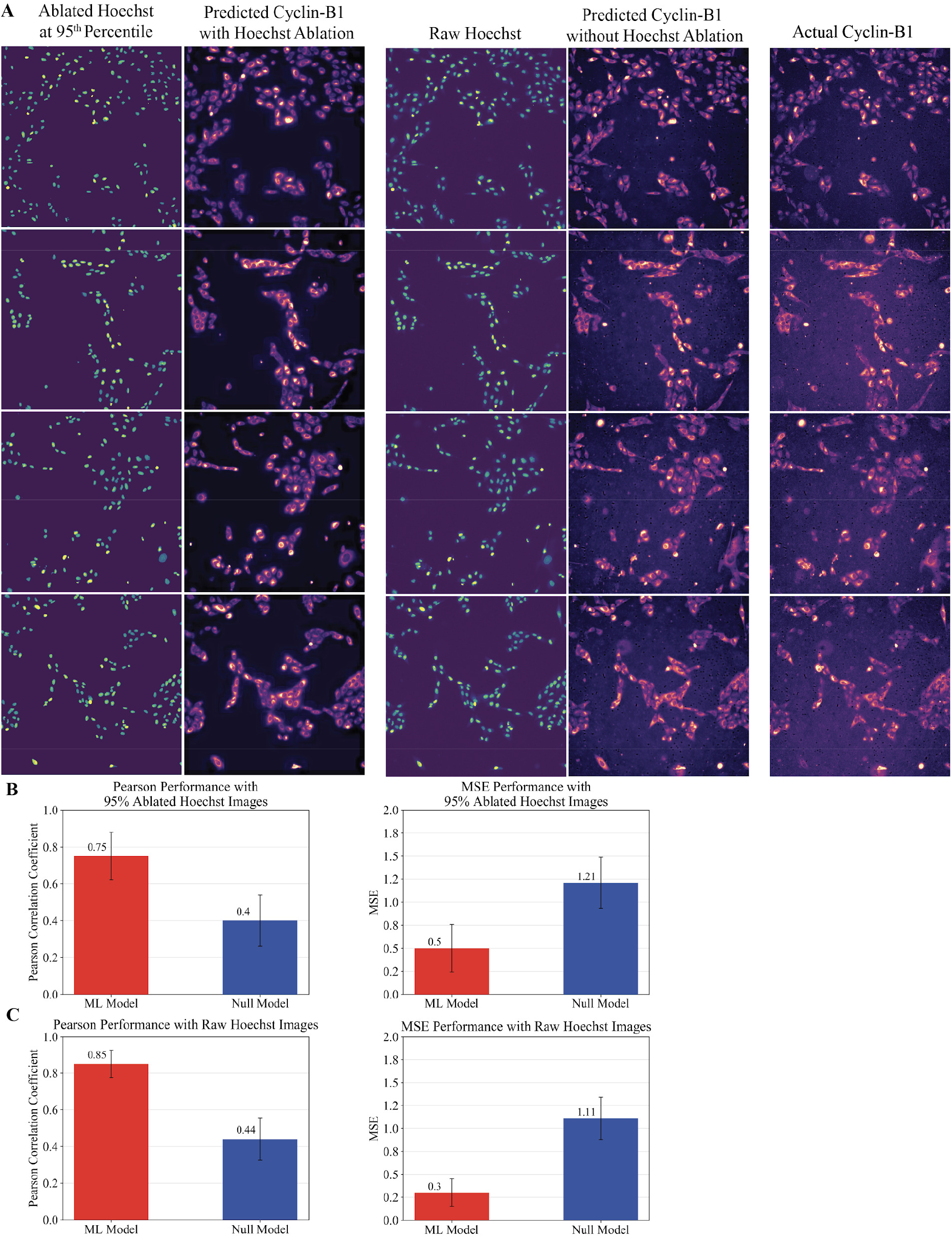
The trans-fluorescence learning method generated accurate cyclin-B1 signal from Hoechst signal, on an independent and biologically unrelated dataset. (a) Given ablated Hoechst images (leftmost column) a ML model trained to output cyclin-B1 images (rightmost column) achieved high *in silico* prediction of this channel (second column). Given unmodified Hoechst stain images (third column), a separate ML model constructed high-fidelity and detailed predictions (fourth column) of the actual cyclin-B1 channel. This panel contains images from the test set (*n*=108,330 images) never seen during model training. For visualization, all images were auto-enhanced and colored as in prior Figures. (b) Assessing the Hoechst-ablated model’s performance. Left: average Pearson correlation coefficient when assessing pixel-wise similarity between actual and predicted Cyclin-B1 images in threefold cross-validation of a dataset of *n*=324,989 total image pairs, across *g*=16,194 unique functional genomic perturbations; Right: MSE from the same analysis. Assessing the ML Model’s superior average performance over the Null Model resulted in statistical significance of p<<0.00001. (c) Assessing the model trained using raw, unablated Hoechst images, performing the same analysis as in (b). The ML Model’s superior average performance over the Null Model likewise resulted in p<<0.00001. Error bars show one standard deviation in each direction centered at the mean.

When we omitted the ablation procedure and trained a threefold cross-validation on raw images, the ML model accurately predicted cyclin-B1 signal and produced detailed and realistic images that matched the observed images to which it had been blinded (Figure 6A). This resulted in an average Pearson correlation of 0.85 ± 0.08 (corresponding to an MSE of 0.30 ± 0.15) on n=108,330 test images (Figure 6C). This was 93.2% higher (p<<0.00001) than the Null Model that stipulated the raw Hoechst image as its prediction (Pearson correlation of 0.44 ± 0.12; MSE of 1.11 ± 0.23). Training on raw, unablated images resulted in a 13.3% (p<<0.00001) increase in average Pearson performance versus the ablated training procedure.

## Discussion

We developed and assessed a method to computationally augment fluorescence microscopy and HCS. This method could be especially useful for tapping into otherwise hidden information in large and information-rich archival datasets. Three observations merit emphasis: (1) trans-fluorescence models generated biologically informative images that were drop-in replacements for existing HCS workflows; (2) the model operated on an archival HCS dataset that was independent from the newly-created training dataset and facilitated discovery of new compounds; (3) the method was generalizable to a different biological environment (a functional genomics screen in an osteosarcoma line).

We readily integrated the ML trans-channel predictions into a conventional compound evaluation workflow because all of the *in silico* channel images were generated as drop-in replacements for the equivalent *in vitro* experiment. Generating a predicted AT8-pTau image from existing YFP-tau and DAPI fusion images required only fractions of a second. With the proper hardware or cloud resources, this method scales to large, high-volume HCS datasets and is parallelizable. Its ability to accurately predict trans-channel fluorescence can potentially serve as a biologically informed and economically practical means of hypothesis generation, especially if the desired marker in question has barriers to widespread utilization. This method is also pragmatic whereinformation-rich bright-field images are not available^32^, which was the case for our screen. Furthermore, accurate *in silico* labeling decreases variability from possible experimental labeling artifacts.

As measured by dose-response confirmed compounds, the ML method performed similarly to the conventional method in the drug discovery task. We were intrigued, however, that the ML method found a comparable number of active compounds, many of which were missed in the original screening campaign. A strong comparison could not be made between the approaches and their propensity to find active compounds at the small scale of 17 unique active compounds, when the methods’ hit rates differed by one hit. Rather, the demonstrated utility of the ML method was to triage compounds more efficiently and to rescue efficacious compounds that were missed due to poor rank in the primary screen. Encapsulating information on AT8-marked phosphorylation and projecting it onto the HCS dataset facilitated compound ranking and subsequent validation.

We acquired the three-channel fixed-cell HCS training dataset using a slightly different protocol than the archival live-cell HCS dataset. Separated by differences in protocol, cell format, experimenters, and a year, the ML model relied solely on extracting hidden signal of phenotypic phosphorylation from YFP-tau images—with no AT8 labeling in the HCS, and thus no possibility of fluorescent bleed-through. This implies that the model has value outside of the strict data regime on which it was trained. This is not always the case in ML applications, which often do not generalize^40^. The archival HCS test dataset consisted completely of compounds that were not part of the training dataset, yet the model was able to operate on new images and a new and larger compound space of greater than a thousand compounds to facilitate discovery.

By evaluating the method on a biologically unrelated osteosarcoma dataset, we showed that the method of trans-channel fluorescent learning generalized across disparate data domains. Intended only as a test in a different disease and biological domain, we did not seek to address nuanced biological questions related to accurately constructing cyclin-B1 signal solely from Hoechst signal. We hope that researchers can tailor the method to their own use cases in order to augment their phenotypic screening endeavors and expand to use cases we have not envisioned. Furthermore, the architecture is fully convolutional, invariant to image size, and adaptable to any input shape. Hence, we expect this method to aid development efforts across a wide range of image applications.

Several caveats limit and focus the scope of this study. The cell line for the tau experiment was not neuronal, and HEK cells were chosen for practicality—HEK cell lines are frequently used in drug screens since they are easy to grow and transfect to express a protein of interest. Additionally, the method, as with most ML applications, was data intensive and required many image examples for learning. This could be a pragmatic hurdle, but we hope to leverage transfer learning^41^ as openly shared cellular image data become increasingly common^42^. Accordingly, we have made all of the tauopathy training dataset and one full plate of the osteosarcoma functional genomics HCS dataset publicly available (Methods). Furthermore, the AT8-pTau channel images collected experimentally could be missing information on tau that is phosphorylated at other epitopes of interest. Thus, what we currently interpret as YFP off-target signal—i.e., the “false positive” pixels that show fluorescence in the YFP-tau but *not* in the AT8-pTau channels—could represent disease-relevant phosphorylation at sites other than the Ser202/Thr205 epitope. A future exercise would be to use antibodies for other commonly hyperphosphorylated epitopes. Consequently, we did not seek with this study to implicate tau hyperphosphorylation as a disease-inducing mechanism, but rather used the AT8 marker as an approximate surrogate for a relevant disease phenotype.

Trans-channel learning is a tool for hypothesis-guided biological discovery. When a new biologically-informative channel can be reliably predicted on an archival dataset, it decodes actionable high-content signal that can guide compound prioritization and rescue missed opportunities hidden in completed HCSs for drug discovery. Conversely, when employed instead as a means to *falsify* hypotheses about the relatedness of biological processes, the failure of a best-effort trans-channel approach to learn one channel when it succeeds in learning other trans-channel relationships may be informative when assessing the biological and visual independence of a pair of biological markers (Supplemental Figure 3). We hope that the techniques and datasets we describe and make available here, which may be attempted on any archived high-content screen, will be of broad use to the microscopy, screening, and drug discovery communities.

## Methods

### Tg2541 Mouse Line

The Tg2541 transgenic mouse line expressed the 0N4R isoform of human tau, under control of the murine neuron-specific Thy1.2 genetic promoter^43^. Tg2541 mice were originally generated on a mixed C57BL/6J × CBA/Ca background^43^ and were then bred onto a C57BL/6J background using marker-assisted backcrossing for eight generations before intercrossing to generate Tg2541 homozygous congenic mice. These mice were kept at room temperature. Homozygous Tg2541 mice on a congenic C57BL/6J background were kindly provided by Dr. Michel Goedert (Medical Research Council, Cambridge, UK). The mice were maintained in a facility accredited by the Association for Assessment and Accreditation of Laboratory Animal Care International in accordance with the Guide for the Care and Use of Laboratory Animals. All procedures for animal use were approved by the University of California, San Francisco’s Institutional Animal Care and Use Committee.

### Phosphotungstic acid precipitation of tau protein in Tg2541 mouse brain samples

Pooled brain homogenate from 6−7 month old Tg2541 mice (both male and female) was prepared as previously described^25^. Briefly, a 10% (weight/volume) homogenate in DPBS was prepared using a rotor-stator type tissue homogenizer (Omni International). Phosphotungstic acid (PTA) precipitation of the brain homogenate was then performed as previously described^12,44^. Briefly, 10% brain homogenate was incubated with 2% sarkosyl (Sigma Aldrich) and 0.5% benzonase (Sigma Aldrich) at 37°C with constant agitation at 1,200 rotations per minute for 2 hours on an orbital shaker. Sodium PTA was dissolved in water and the pH was adjusted to 7.0. A final concentration of 2% sodium PTA was added to the brain homogenate and incubated overnight under the same agitation conditions. The brain homogenate was then centrifuged at 16,100 × g for 30 minutes at room temperature, and the supernatant was removed. The pellet was resuspended in 2% sarkosyl and 2% sodium PTA in DPBS and incubated for 1 hour at room temperature. The sample was centrifuged again under the same conditions, the supernatant was removed, and the pellet was resuspended in DPBS, using 10% of the initial starting volume, and stored at −80°C until further use.

### Experimental Setup for the Cellular Tau(P301S)-YFP Aggregation Assay Training Dataset

Tau(P301S)-YFP cells were developed by transfecting human embryonic kidney cells (HEK293T female; ATCC) using electroporation to overexpress the full-length 0N4R isoform of human tau containing the familial disease-linked P301S missense mutation and the yellow fluorescent protein (YFP) fused to the C-terminus for visualization. A stable monoclonal line was maintained in Dulbecco’s modified enriched medium (DMEM) supplemented with 10% fetal bovine serum (FBS) and 1% penicillin/streptomycin. Tau(P301S)-YFP cells in confluent flasks were collected using trypsin and resuspended in electroporation buffer at a concentration of 8.4 × 10^7^ cells/mL. HEK293T cells that overexpress the microtubule-binding repeat domain (RD) of 4R human tau with the P301L and V337M mutations^13^ [TauRD(P301L/V337M)-YFP cells] were maintained in DMEM supplemented with 10% FBS and 1% penicillin/streptomycin. Tau seeds were prepared by passaging PTA-precipitated tau protein from Tg2541 mouse brain samples once through TauRD(P301L/V337M)-YFP cells and then collecting the cell lysate, as described previously^25^. All cells were cultured at 37°C and 5% CO_2_ in a humidified incubator until analysis.

Cell lysate containing tau seeds was combined with the Tau(P301S)-YFP cells at 1.8 µg/mL and electroporation was performed on an Amaxa Nucleofactor instrument (Lonza Bioscience) using the program #Q-001. Seeded Tau(P301S)-YFP cells were then plated at 5 × 10^3^ cells/well in a 384-well plate (Greiner) coated with poly-L-ornithine (PLO, 30-70kDa; Sigma Aldrich). To coat plates, PLO was dissolved at 1 mg/mL in 0.15 M sodium borate buffer pH 8.4 and filter-sterilized, and then further diluted to 0.1 mg/ml in sterile deionized (DI) water. The PLO coating solution was added to the culture plate wells and incubated in culture plate wells for six hours at room temperature, then removed. The wells were washed three times with sterile DI water and air-dried in a biosafety cabinet.

Immediately following cell plating, small-molecule drug compounds prepared in pure DMSO were added at a fixed concentration of 10μM. The small molecules were selected to represent the range of effects on tau aggregation and cell viability by compounds in drug libraries of interest, developed by Daiichi Sankyo (Tokyo, Japan). Actual compound identifiers are proprietary and are obfuscated to identifiers in the form “DRW#”. Two compounds were selected for reducing tau aggregation and also reducing cell viability (DRW1 and DRW2); and two compounds were selected for increasing tau aggregation and reducing cell viability (DRW3 and DRW4); two compounds were selected for reducing intracellular tau aggregation without affecting cell viability (DRW5 and DRW6). Control wells were also prepared with cells electroporated with or without tau seeds and the same volume of DMSO but no drug compound.

At four days post-seeding, cells were washed three times with sterile Dulbecco’s Phosphate (DPBS) and then fixed with 4% paraformaldehyde (PFA) in DPBS for 20 minutes at room temperature. Cells were washed with DBPS and then permeabilized with 0.1% Triton X-100 in DPBS for 20 minutes at room temperature. Cells were washed and then blocked with 3% bovine serum albumin (BSA; Millipore) in DPBS for one hour at room temperature. Cells were then incubated with mouse monoclonal antibody AT8 (1:500; Thermo Fisher Scientific) and rabbit polyclonal antibody MCM2 (1:500; Abcam) in 3% BSA in DBPS overnight at 4°C. Cells were washed with DBPS and then incubated with Alexa Fluor 594 Plus-conjugated goat anti-mouse and anti-rabbit secondary antibodies (1:500; Thermo Fisher Scientific) in 3% BSA in DBPS for 90 min at room temperature. Cells were washed with DBPS and then covered with DPBS and imaged on an InCell Analyzer 6000 High-Content Confocal Microscopy System (GE Healthcare). Twenty five non-overlapping, 2048 × 2048 pixel fields were captured per well. We chose markers with minimal overlap of the excitation spectra to minimize potential image bleed-through (Supplemental Figure 6A). This full training dataset was uploaded here for open access (DOI: 10.17605/OSF.IO/XNTD6).

### Constructing the Archival HCS Dataset

The experimental setup for the archival HCS data was the same setup used to construct the training set (the cellular Tau(P301S)-YFP aggregation assay), with the exceptions that the HCS data did not undergo any of the immunocytochemistry steps, and live-cell imaging was used instead. We constructed this HCS dataset more than one year prior to the three-channel training dataset. Only two imaging channels were captured, a nuclear DAPI channel and a YFP-fused tau channel. No AT8 antibody was plated. Thus no AT8-pTau channel was captured, and there was no possibility of bleed-through. Furthermore, the compound library in the HCS data was much larger and more diverse than that used for model training. The original six compounds used in the training set with AT8 were not included in the random HCS subset of 1,600 unique compounds evaluated by the ML model.

### Training and Evaluation of the ML Models from the Tauopathy Dataset

The full source code and fully trained models are available at https://github.com/keiserlab/trans-channel-paper. The necessary python packages are PyTorch, pandas, cv2, cupy, numpy, and sklearn. Training and testing of the model was performed on one Nvidia Geforce GTX 1080 graphics card, and completed in 30 epochs totaling 111 hours of training.

The full dataset consisted of six different drug perturbations: two reduced aggregation while maintaining cell count; two reduced aggregation while reducing cell count; and two increased aggregation while reducing cell count. We imaged a total of six 384-well plates, one plate for each drug. Each well was divided into four fields and each field was imaged, resulting in a full image dataset consisting of 57,600 TIFF images each of size 2048 × 2048 pixels for each of the three channels. As a preprocessing step, we linearly scaled all images from the original TIFF range of 0 to 65,535 to the more tractable range of 0 to 255 by dividing each pixel by 65,535 and then multiplying by 255. We converted the resulting images to 32-bit floating point. The images were randomly shuffled, and then split into a 70% training set and 30% test set.

We used the fully convolutional architecture in Figure 2A for training. As inputs to the model, we concatenated the transformed YFP-tau image and the transformed DAPI image into a tensor of size 2 × 2048 × 2048 pixels. The model generated a predicted AT8-pTau image of size 2048 × 2048 pixels.

We used a stochastic gradient descent optimizer with momentum = 0.90, a learning rate = 0.001, and a batch size of one. We trained the model for 30 epochs. We used a negative Pearson correlation loss function for training, in which we minimized the following objective function:

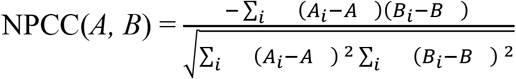

such that *A* and *B* were the images being compared, *i* was summed over all pixels,

A_μ_ was the average of image A, and B_μ_ was the average of image B

### Novel Model Design Considerations

Our architecture differed from U-Net in several important ways (Supplemental Figure 1). First, the model was tasked with assigning pixel values that resided in a greater range (0 to 65,535) compared to U-Net’s binary zero or one objective. Hence we used a negative Pearson correlation loss while U-Net used a cross-entropy loss. Our architecture had fewer hidden layers than U-Net, resulting in an architecture that was 12% the size of U-Net. Also the upsampling procedure was different. We used transposed convolutions followed by bilinear interpolation, while U-Net had two convolutions followed by bilinear interpolation. We used transposed convolutions for learnable upsampling during each convolution operation. Finally, our architecture preserved the original pixel image dimensions, while U-Net returned a smaller mask. Preserving image dimensions allowed for a direct pixel-wise one-to-one mapping from the input to the output.

### Performance Metrics

When constructing the Pearson performance metric, we iterated over the test set, flattened all of the images to one dimensional vectors, and then found the Pearson correlation of the flattened label image with the flattened ML-predicted image using the Numpy library’s corrcoef function.

When constructing the MSE metric, we normalized all AT8-pTau and predicted AT8-pTau images to have zero-mean and unit-variance in order to account for differences in underlying pixel value distributions between the label images and the predicted images. Normalizing both distributions placed them in a more comparable regime for calculating MSE.

When calculating MSE, we normalized each image by subtracting the average of that image, and then dividing by the standard deviation. The MSE was determined as follows:

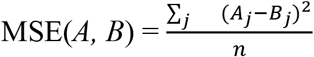

Such that *A*_*j*_ and *B*_*j*_ were the normalized images being compared, and *j* was the index of the image out of a total of *n* test images.

To construct the ROC curve, we first normalized the images to have zero-mean and unit-variance. We then compared the predicted image to the actual, and chose a pixel threshold *t* such that any pixel greater than or equal to *t* was considered as positive for signal, and anything lower than *t* was negative for signal. We chose an intensity threshold of 1.0 to binarize the image, which retained most of the features of interest and aggregation signal (Supplemental Figure 9).

Hence, we performed pixel wise classification, and derived an ROC curve for each image. The ROC curve in Figure 3 is the average ROC curve across the full test set of images. For the thresholds applied to the predicted image, we chose the pixel value range 0 (permissive) to 1000 (non-permissive) with different increments. In the range -1.0 to -0.1 and the range 0.1 to 1.0, we chose incrementing values of 0.1. From the range -0.1 to 0.1 we chose a more fine grained increment with a step size of 0.01. In the range 1.0 to 4.0 we chose a step size of 1.0 because of the infrequency of these values. Finally, we evaluated the pixel intensity threshold of 1,000, for which no pixels assumed this value.

### Construction and Evaluation of PQML and PQC

From the available YFP-tau fusion images and DAPI images of the HCS subset of 1,600 compounds, we constructed predicted AT8-pTau images for each pair of YFP-tau and DAPI images. We took the YFP and DAPI images and linearly scaled the pixel values to the range 0 to 255 inclusive. We then concatenated the two images together to form a tensor of dimensions 2 × 2048 × 2048, and finally inputted this into the trained model. Next, we scaled the output of the model back to the original space by first dividing by 255 and then multiplying by the maximal possible pixel value of 65,535. We then scored these images for tau aggregation using proprietary software from General Electric, called the “InCell Analyzer,” which yielded a score based on puncta count and area. This same algorithm was used for previous HCS efforts, and it was applied consistently to all images. We obtained each small molecule’s aggregation score by averaging over all of that compound’s images’ aggregation scores—which consisted of averaging over four non-overlapping fields within a single well treated with the compound. We ranked all of the compounds by their aggregation score, with compounds inducing lower aggregation being higher priority than compounds with a higher aggregation score. A lower rank in the queue indicated higher priority (e.g. compounds with ranks first, second, and third were the three highest priority compounds). For constructing PQC, we used the same method as the one to construct PQML, except that we obtained aggregation scores from the original YFP-tau images instead of their corresponding ML-predicted AT8-pTau images.

To calculate enrichment curve AUCs (Figure 5), we analyzed the set of known active compounds (*n*=17) discovered in the study. For each queue, we generated an enrichment curve using the rankings of the active compounds. We calculated AUC by integrating with Numpy’s composite trapezoidal function (numpy.trapz). We calculated average active compound ranks by averaging over each active compound’s index in the queue (Figure 5B).

### Experimental Setup for the Secondary Dose-Response Experiments

For the dose-response experiments, we performed the same protocol as the cellular Tau(P301S)-YFP aggregation assay, with the exceptions being the drug doses and the compounds that we tested. We plated drug concentrations in half logs from 10nM to 10μM. We tested the top 40-ranked compounds of PQML and the top 40-ranked compounds of PQC. In accordance with the cellular Tau(P301S)-YFP aggregation assay, we did not use AT8 because it would demand significant additional time and labor. For each compound, we independently replicated each concentration in four separate wells. We calculated the average aggregation scores at each concentration, and used these averages to construct a dose-response plot (e.g., Figure 4A).

### Assessing Activity from Dose Response Curves

A compound that succeeded in a secondary dose-response test demonstrated a dose-dependent decrease in aggregation as a function of concentration, while maintaining a therapeutic window that preceded a noticeable drop in cell count. A compound did not need to induce a perfect sigmoidal shape to be considered active. A compound was considered active if and only if all of the following four conditions were satisfied.

1. The best fit curve strictly had an aggregation score that decreased with increasing concentration of compound.
2. The effective response was at least 6,000 aggregation units. We required this so that only compounds were chosen that induced an effective decrease in aggregation. Medicinal chemists who worked closely on the screen chose 6,000 subjectively as a minimal threshold before we began the analyses. If the curve was monotonically increasing with increasing concentration, this condition was automatically not satisfied, and the compound was considered inactive.
3. The half maximal effective concentration (EC50) was lower than 10μM.
4. The concentration at which aggregation began to be ameliorated at half maximal response was lower than the concentration at which the compound decreased cell count by half. If the cell count trend increased with concentration, or was unaffected, then this condition was automatically satisfied. Otherwise, the logarithm of the EC50 of the aggregation curve minus the logarithm of the EC50 of the cell count curve must be less than -0.10. We chose -0.10 as a more stringent threshold, as opposed to the difference being anything less than 0.0. This ensured that the compound’s concentration at which it exhibited half of its maximum response occurred at a lower concentration than when the compound began to decrease cell count by half, and that this difference was not effectively zero.

### Experimental Setup for Validation Assay of Functional Genomics Screen in Osteosarcoma Line

We generated a completely different HCS experiment than the tauopathy study in order to assess the ML method’s generalizability. This osteosarcoma dataset was subjected to completely different biological conditions. U2OS cells (female) expressing a stable CCNB1-GFP construct were plated into 384 well plates with 500 cells per well and reverse transfected with an esiRNA library (10ng, Sigma Aldrich) using Hiperfect transfection reagent. The U2OS cells were cultured in DMEM medium containing 10% foetal bovine serum. The library can be found at https://iccb.med.harvard.edu/sigma-esirna-human-1 and also https://iccb.med.harvard.edu/sigma-esirna-human-2. Our method was inspired by the assay presentented in Bray et al.^45^ We performed 16,194 unique functional genomic perturbations. After 72 hours, cells were stained with 5μg/mL of Hoescht 33342 dye (ThermoFisher), incubated for 60 minutes at 37°C, washed with PBS, fixed in 4% PF, and scanned on a Thermo Cell Insight NXT high content microscopy system using a 10X objective. We captured the two channels, resulting in 324,989 Hoechst and cyclin-B1 image pairs, each of dimension 1104 × 1104 pixels.

### Training and Evaluation of the ML Models Used for the Osteosarcoma Dataset

We performed two training experiments, one using ablated Hoechst images, and one using the raw Hoechst images. For the training using ablated Hoechst images, we ablated our Hoechst images at the 95^th^ percentile pixel-intensity threshold. Afterwards, we performed the same exact training procedure as the one we used for the tauopathy experiment, except for the following exceptions: 1) the input to the model was one dimensional (the Hoechst channel) instead of two; 2) We trained for 10 epochs instead of 30 epochs; 3) We trained a threefold cross-validation instead of a single random training-test split.

For training using the unablated, raw images, we applied the same preprocessing and training procedure as the one used for training with ablated images, except for the following exceptions: 1) we left the images intact and did not perform any ablations; 2) we trained for 5 epochs instead of 10. We trained for a shorter time because the model converged faster, likely because the task of learning from raw unablated images was easier.

### Quantification and Statistical Analyses

For all statistical significance tests, we used a two sample, one-sided z test. The null hypothesis stated that performance means were equal. The alternative hypothesis stated that the average performance from ML is greater than the average performance from the non-ML approach. Significance was set at p < 0.05. The values of *n* correspond to the number of instances in the test set (see Method Details for values of *n*). Sample sizes were sufficiently large to perform the test. A sample is one comparison between a predicted channel and the actual channel. We assume normality due to the large sample size.

### Key Resources Table

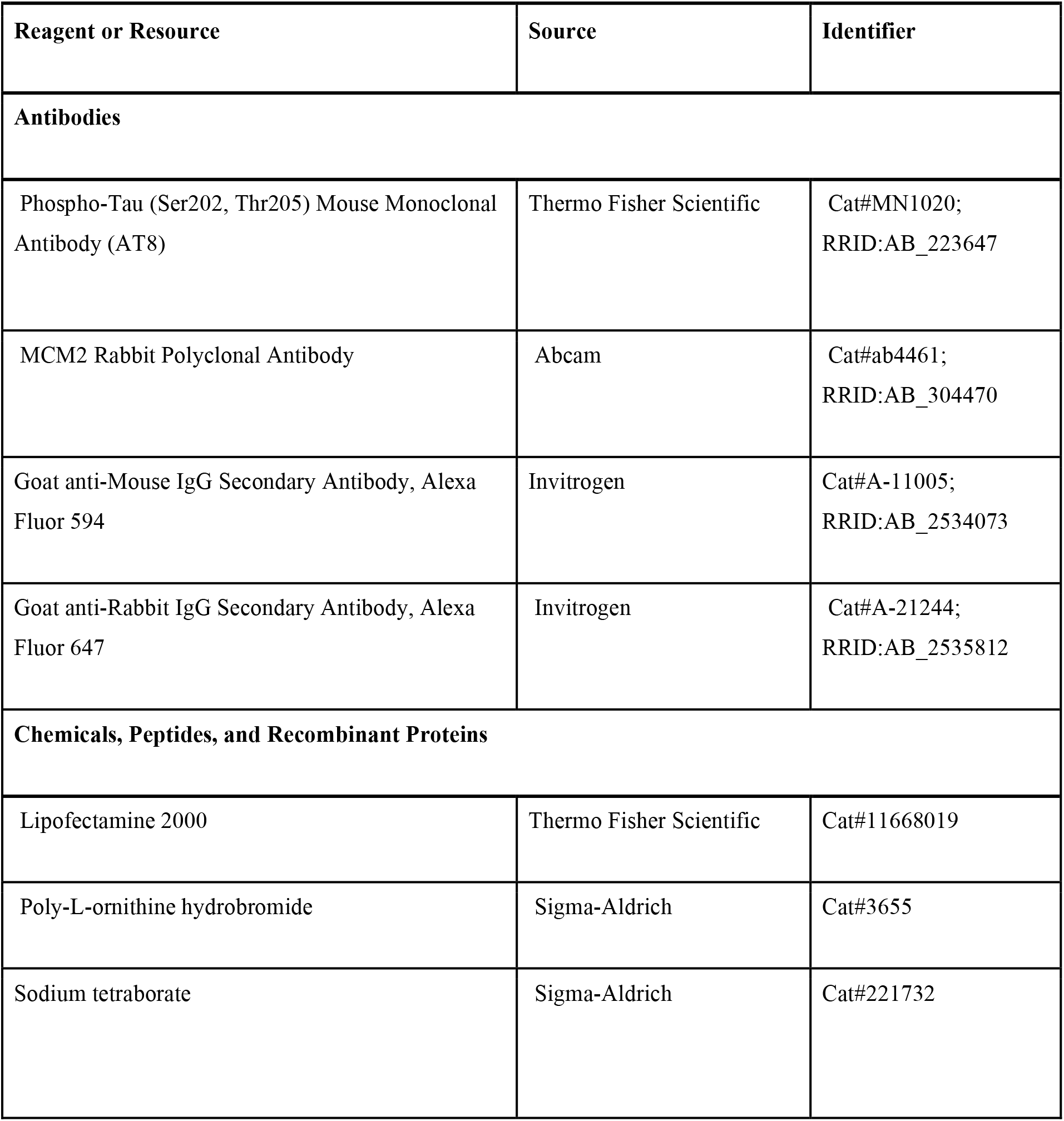
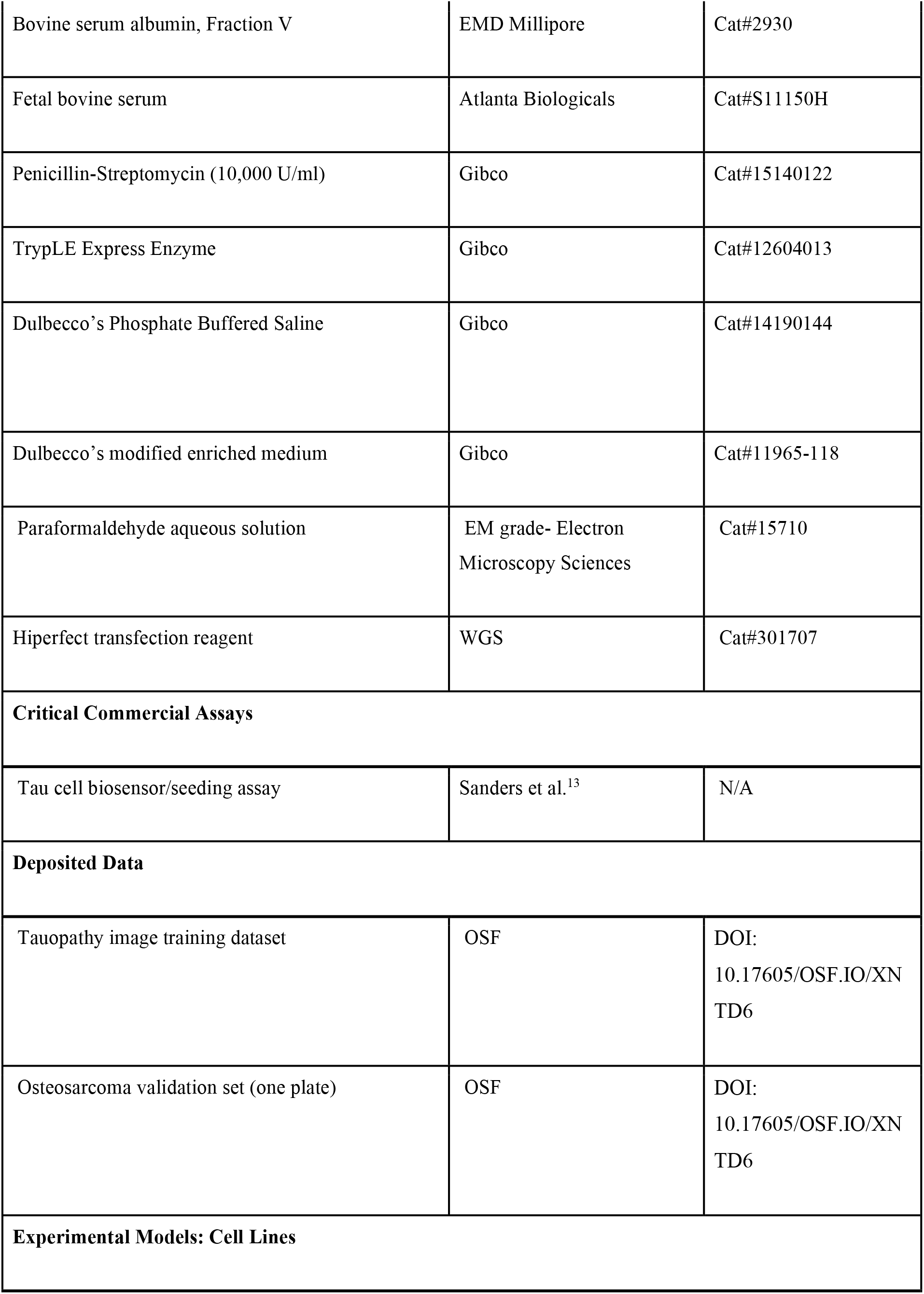
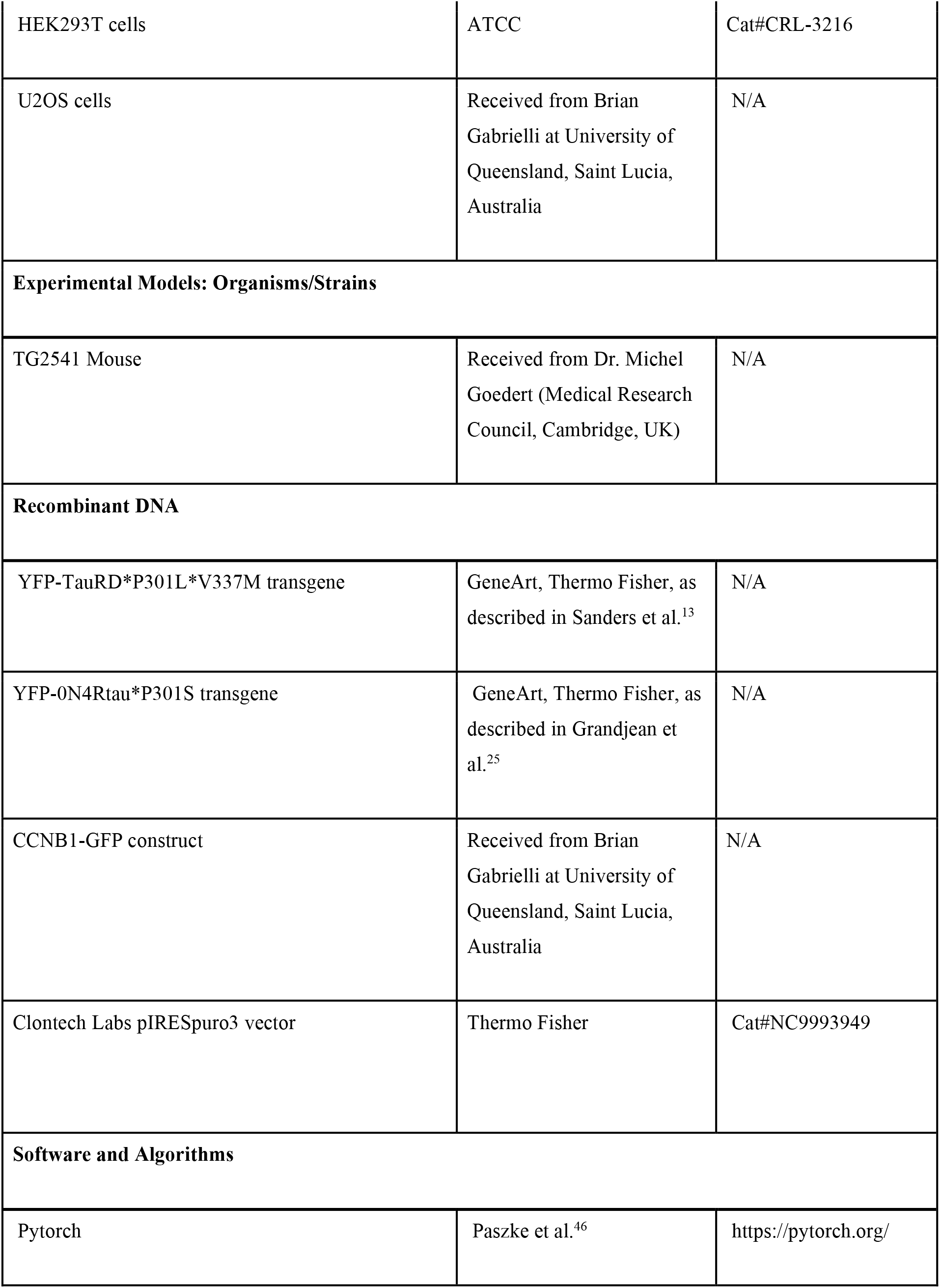
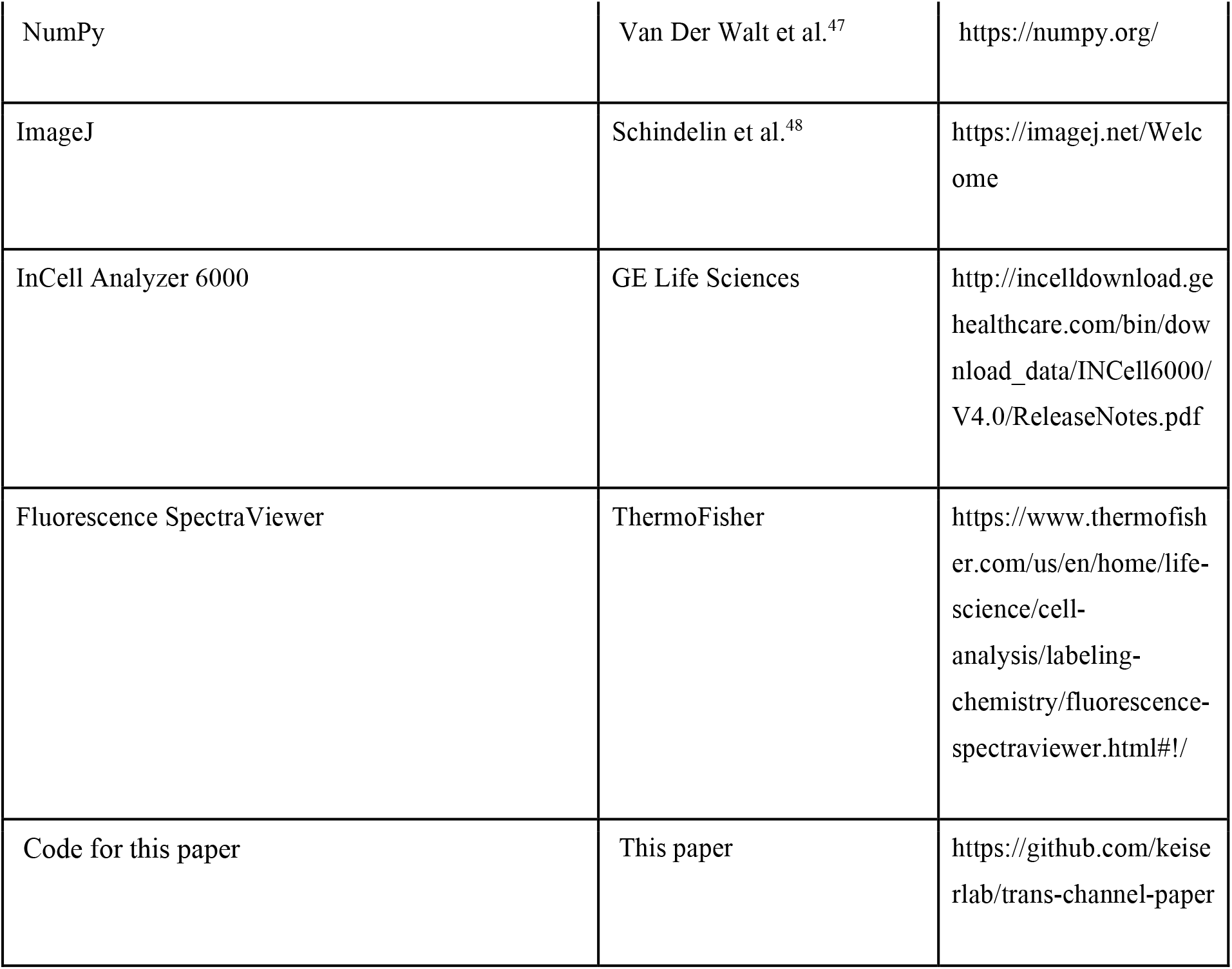

## Author Contributions

Conceptualization, D.R.W., J.C., N.J., N.P., and M.J.K.; Methodology, D.R.W, J.C., N.J., N.P., and M.J.K; Software, D.R.W.; Validation D.R.W., A.L., and S.B.P.; Formal Analysis, D.R.W; Investigation, D.R.W, J.C., N.J., J.C.L, J.A., J.L., and A.L.; Resources, S.B.P, S.B., A.J.B, N.P., and M.J.K; Data Curation, D.R.W; Writing – Original Draft, D.R.W.; Writing – Review & Editing, D.R.W., and M.J.K; Visualization, D.R.W.; Supervision, S.B.P, S.B., A.J.B, N.P., and M.J.K.; Project Administration, D.R.W., and M.J.K.; Funding Acquisition, S.B.P, and M.J.K.

## Acknowledgements

This work was supported by grant number 2018-191905 from the Chan Zuckerberg Initiative DAF, an advised fund of the Silicon Valley Community Foundation (MJK), the National Institutes of Health (AG002132) (S.B.P.), as well as by support from the Brockman Foundation (S.B.P.) and the Sherman Fairchild Foundation (S.B.P.).

## Data Availability

All image data is freely available at this DOI: 10.17605/OSF.IO/XNTD6

## Code Availability

The full source code and fully trained models are available at: https://github.com/keiserlab/trans-channel-paper.

