## Supplementary Material for "Trans-channel fluorescence learning improves high-content screening for Alzheimer’s disease therapeutics"

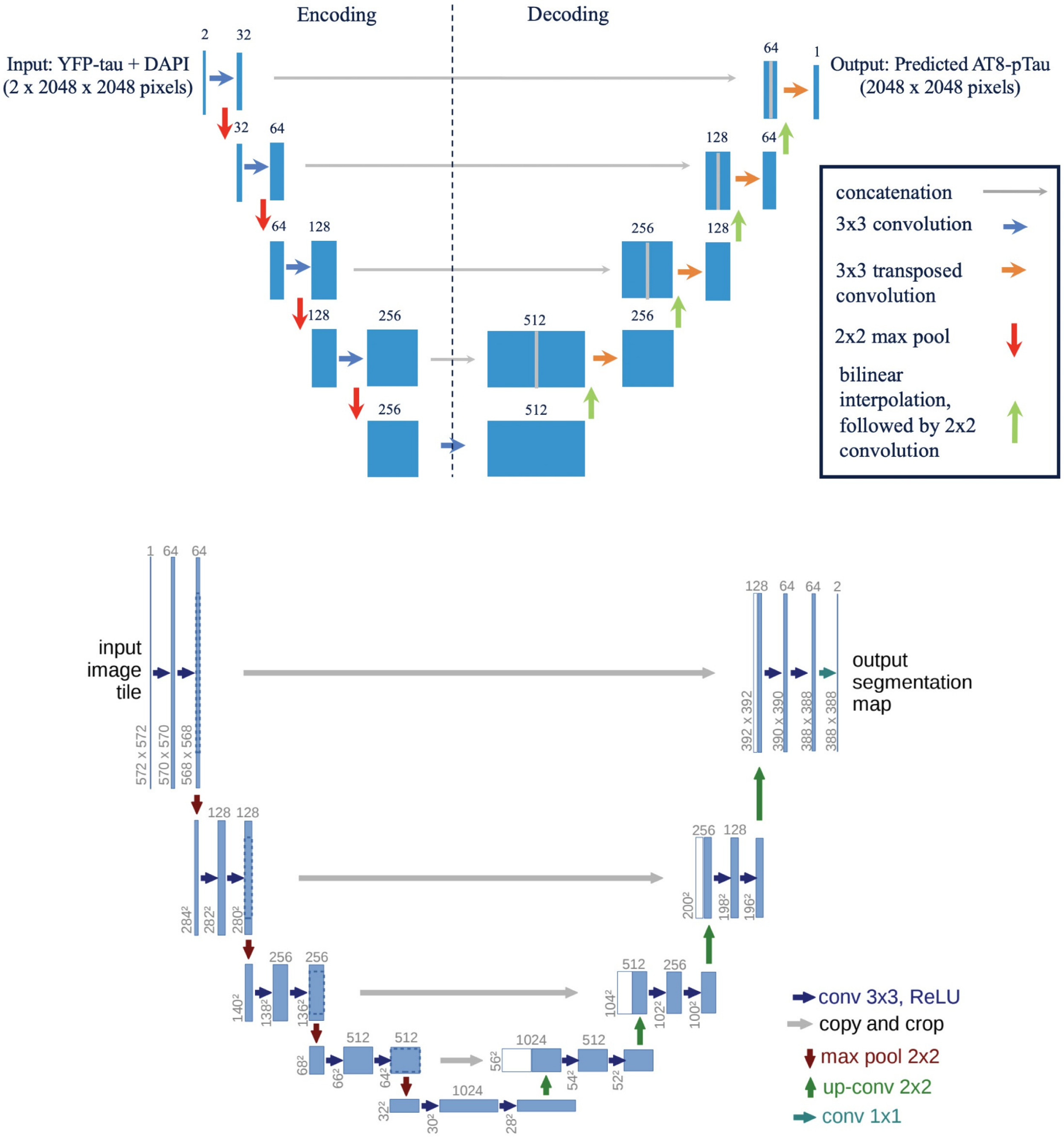


**Supplemental Figure 1: Comparison between our architecture and U-Net.** Top: our architecture, below: the original U-Net architecture. Our smaller, more space-efficient architecture (12% the size of U-Net) preserves image dimensions and yields a model capable of trans-channel learning, with a direct one-to-one correspondence between the input and output image unlike U-Net. Our model learns an image channel that takes on a range of pixel values, in contrast to U-Net which learns a binary mask (see Methods).

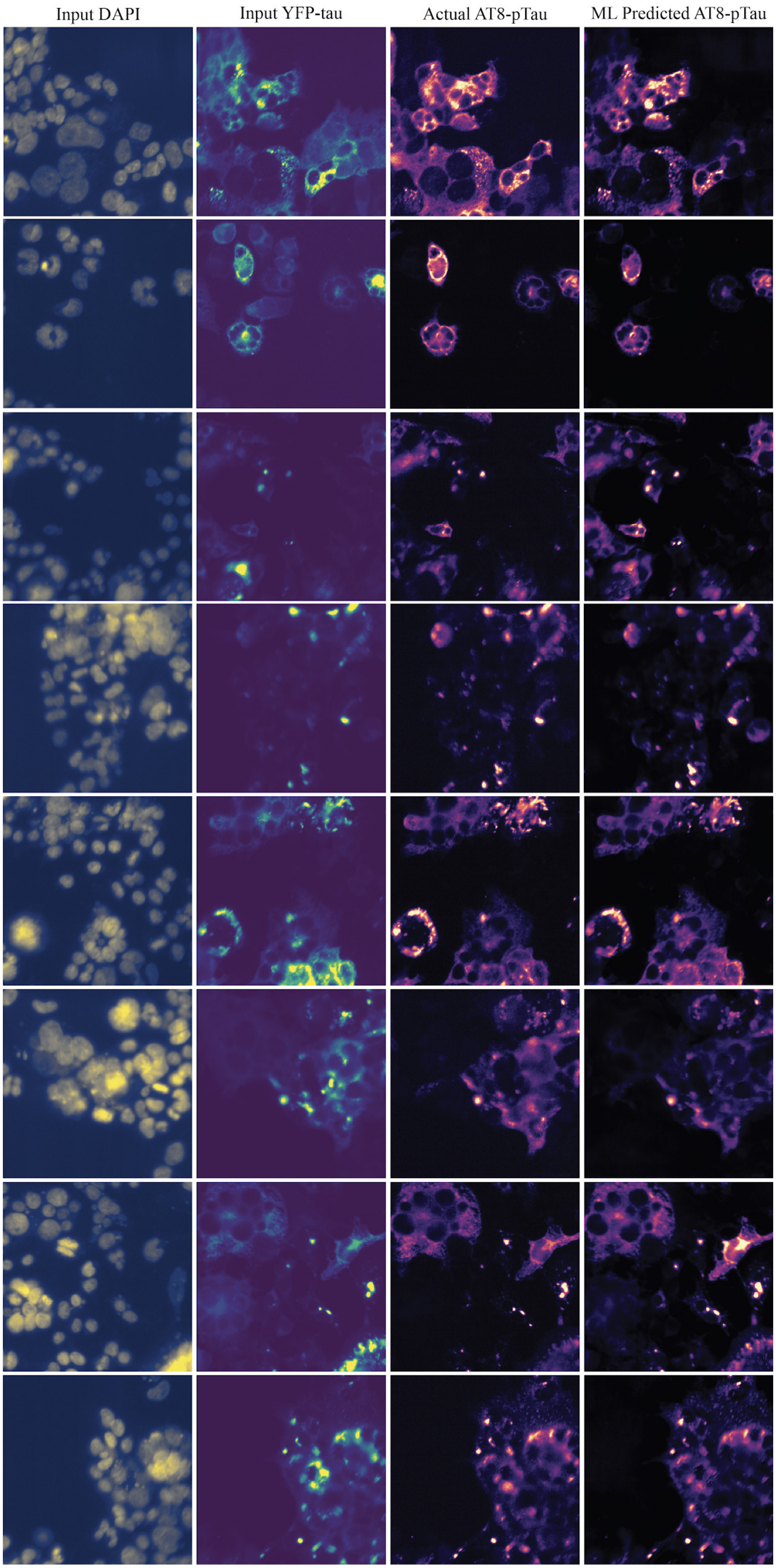
Input DAPI
Input YFP-tau
Actual AT8-pTau
ML Predicted AT8-pTau

**Supplemental Figure 2: Additional image examples showing trans-channel predictions.** The DAPI channel (leftmost) is concatenated with the YFP-tau channel (second column) and inputted to the trained ML Model, which produces a prediction of the AT8-pTau image channel (rightmost). This prediction has high similarity to the actually imaged AT8-pTau channel (third column). All images are pulled from the test set never seen during training.

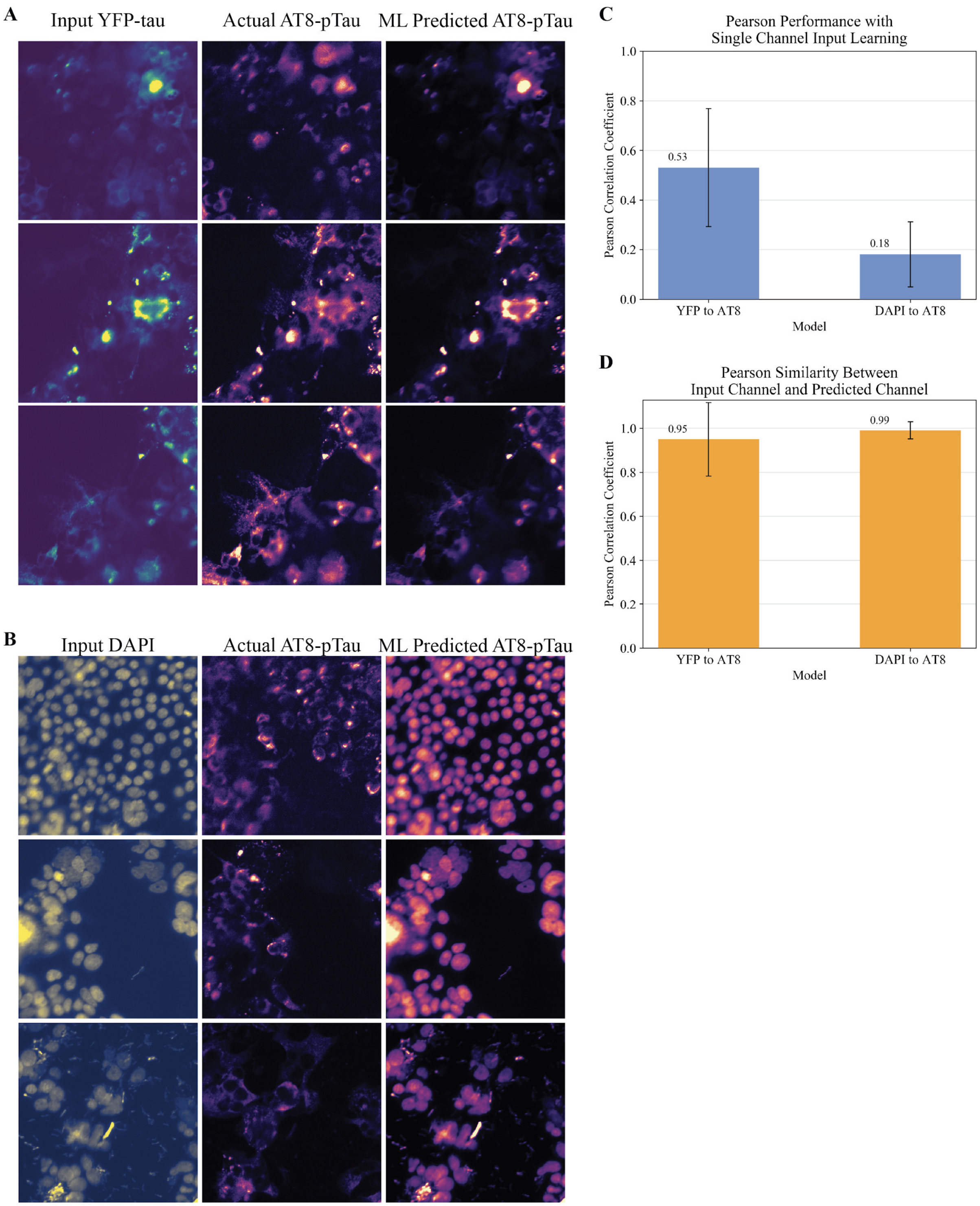


**Supplemental Figure 3: Transchannel-learning from single channel input to objective AT8.** (a) Example input, label, and predicted images from a model trained to map just the YFP-tau channel (without the DAPI channel) to the AT8-pTau channel. (b) Example input, label, and predicted images from a model trained to map just the DAPI channel (without the YFP-tau channel) to the AT8-pTau channel. (c) Pearson performance of models trained with just a single channel, left: model trained to map just YFP-tau to AT8-pTau, right: model trained to map just

DAPI to AT8-pTau. The YFP-tau to AT8-pTau model's performance was nearly identical to the Null Model that simply returned the input YFP-tau channel. The DAPI to AT8-pTau model failed to learn the AT8-pTau channel, as we would expect if the DAPI and AT8-pTau channels are not sufficiently biologically related. (d) Pearson correlation between the single channel model's input and the model's prediction. For both the YFP-tau to AT8-pTau model and the DAPI to AT8-pTau model, the input image is almost identical to the predicted image, indicating that the model learned to simply return its input as the final prediction to maximize the Pearson correlation coefficient. Error bars for (c) and (d) span one standard deviation in each direction, centered at the mean.

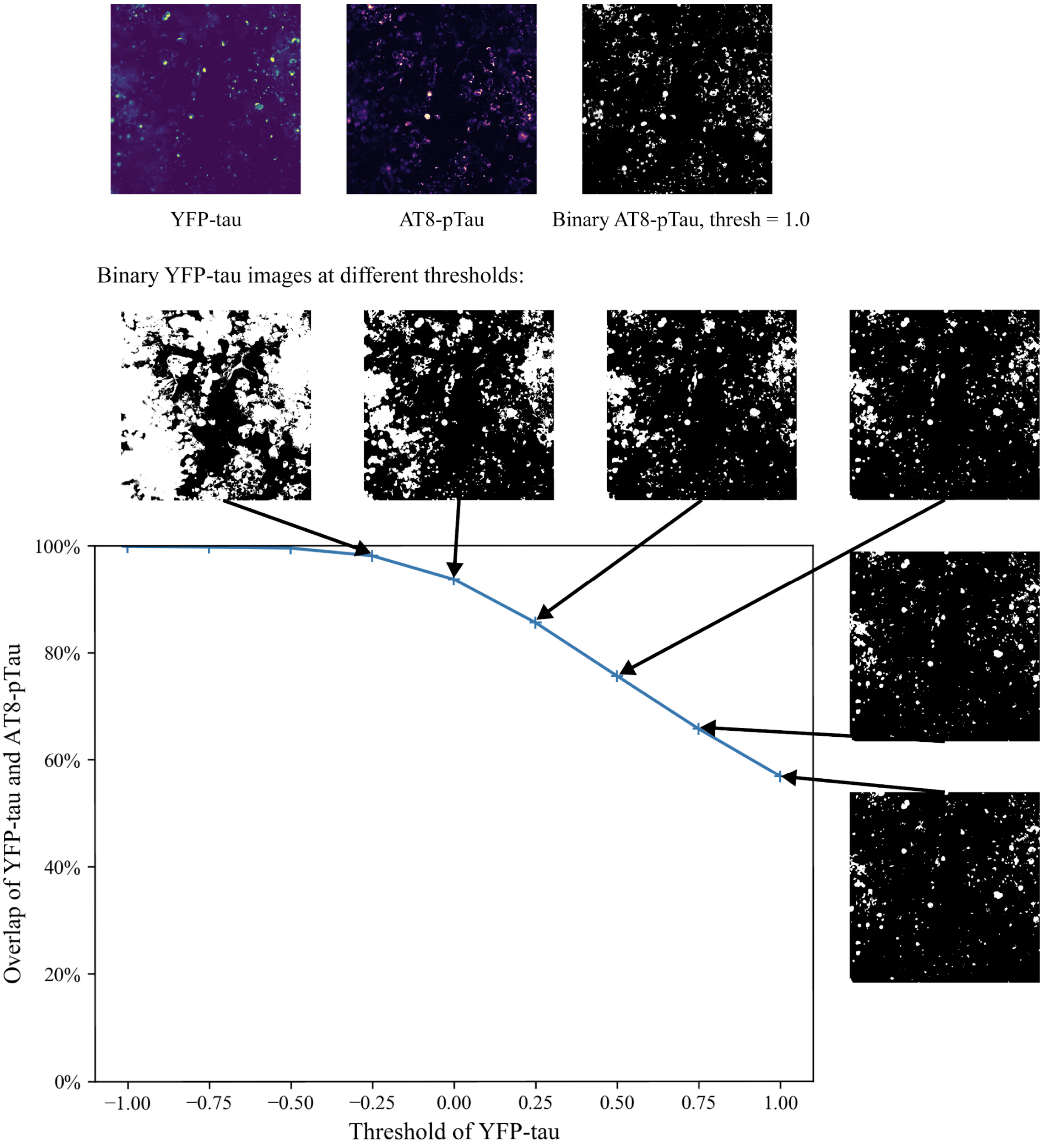


**Supplemental Figure 4: Overlap of AT8-pTau signal and YFP-tau signal.** The x-axis shows the pixel value threshold that binarized each YFP-tau image, such that any pixel greater than or equal to the threshold was considered positive for signal, and anything below this threshold was considered negative for signal. The y-axis shows the average overlap between YFP-tau and AT8-pTau signal (at a threshold of 1.0) as a fraction of AT8-pTau positive signal that was also present in YFP-tau at the same pixel location. We calculated overlaps over the entire image dataset. Example binarized YFP-tau are shown at each threshold. Images at threshold = -0.50 and lower are not shown and consist of mostly signal-positive white images.

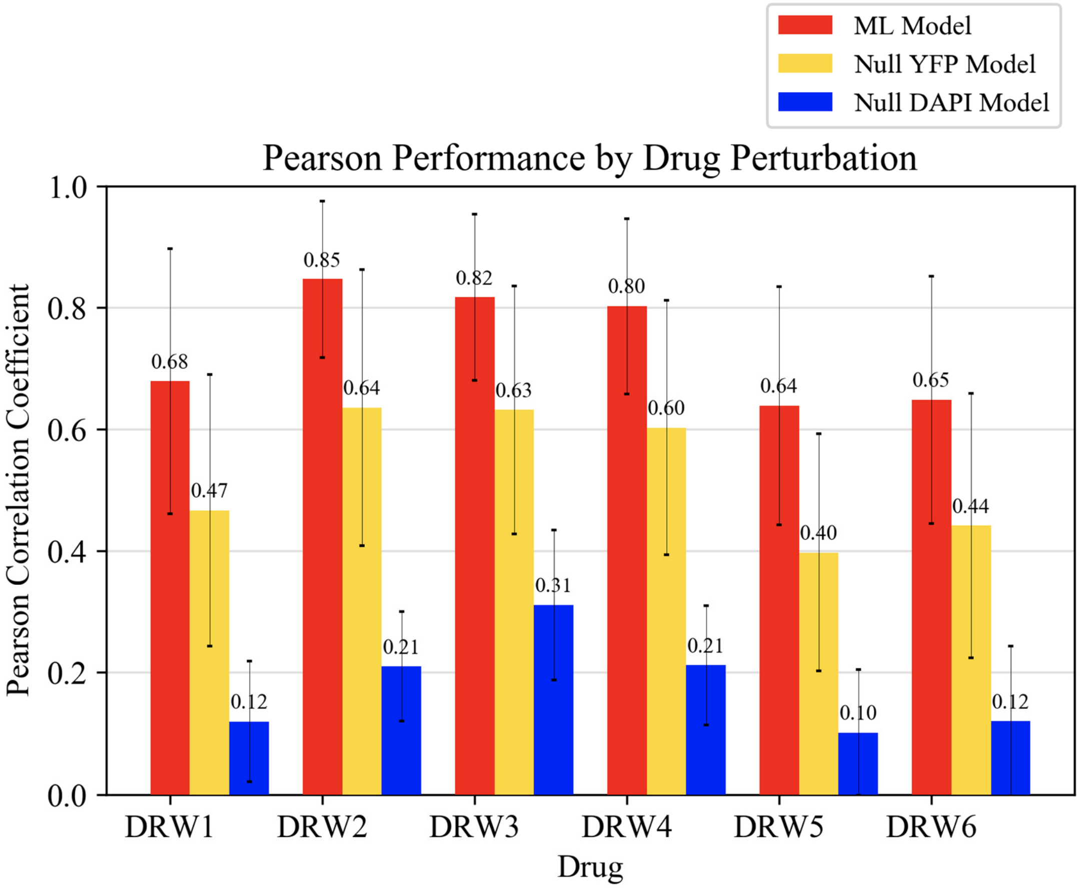


**Supplemental Figure 5: Pearson performance by drug condition.** Performance by drug across the 17,280 held-out test images. Each test image comes from one of six drug perturbations. Error bars show one standard deviation in each direction centered at the mean.

A

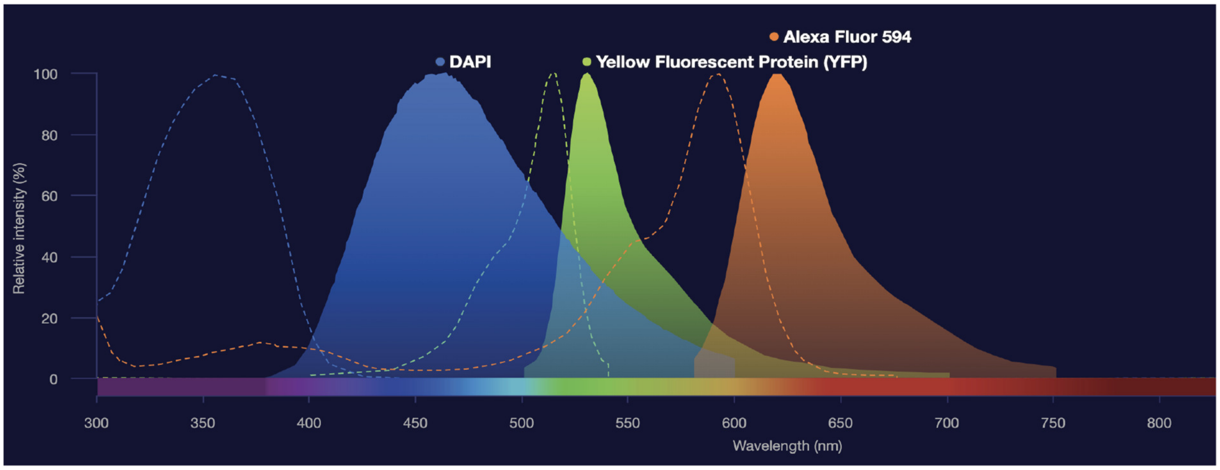


B

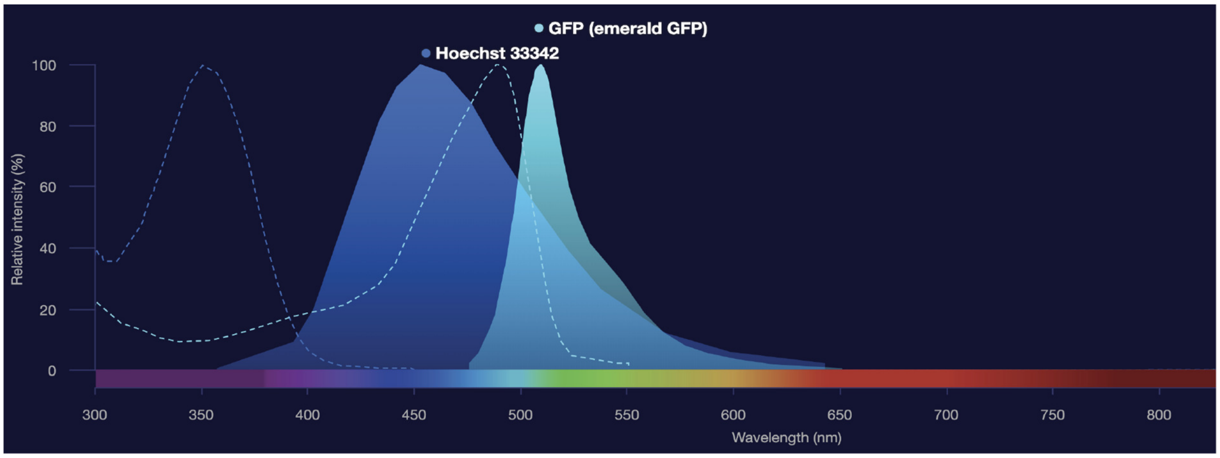


**Supplemental Figure 6: Excitation and emission spectra of fluorophores used in the two biologically distinct datasets.** (a) Excitation and emission of the three-channel tauopathy training dataset. The x-axis plot the wavelength, and the y-axis plots the relative intensity. The dashed lines indicate the excitation plots, and the solid lines indicate the emission plots. Alexa Fluor 594 refers to the secondary antibody used for the AT8-pTau channel (b) Excitation and emission of the functional genomics screen in osteosarcoma cells. These graphs were constructed with ThermoFisher's Fluorescence SpectraViewer: https://www.thermofisher.com/us/en/home/life-science/cell-analysis/labeling-chemistry/fluorescence-spectraviewer.html#!/

**A**

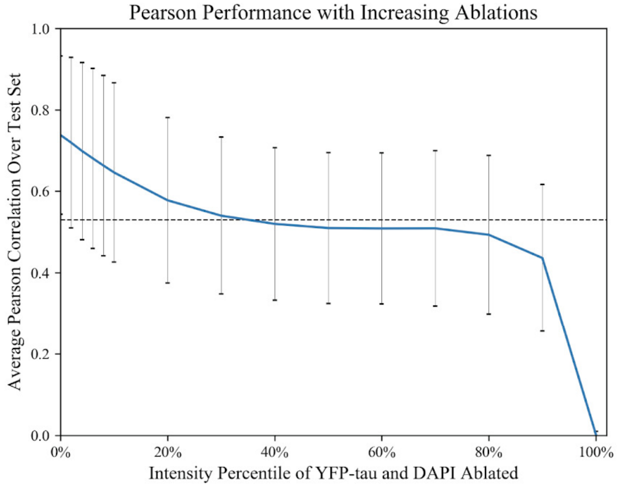
Pearson Performance with Increasing Ablations
Average Pearson Correlation Over Test Set
1.0
0.8
0.6
0.4
0.2
0.0
0%
20%
40%
60%
80%
100%
Intensity Percentile of YFP-tau and DAPI Ablated

**B**

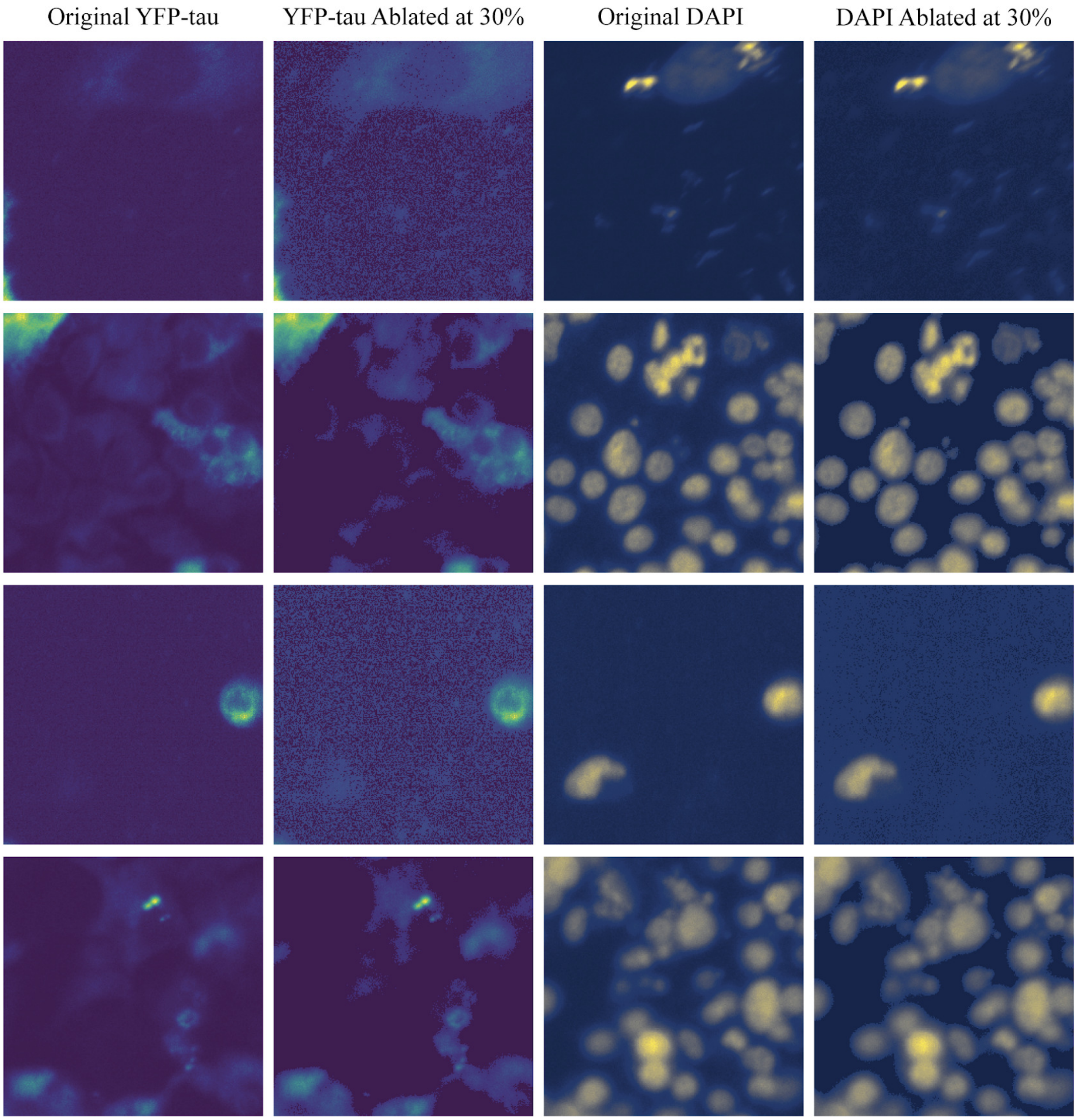
Original YFP-tau
YFP-tau Ablated at 30%
Original DAPI
DAPI Ablated at 30%

**Supplemental Figure 7: Ablation testing on the tau dataset.** (a) We systematically ablated the input channels (both DAPI and YFP-tau) and assessed its effect on performance. We incrementally ablated our input images by reassigning pixels under a specific pixel-value threshold *t* to the value of 0. The threshold *t* was increased from no ablation to a complete erasure of the image. The y-axis plots the average Pearson correlation over the full test set at different percentile ablations (x-axis). We performed a more fine grained search at low ablations where faint bleed-through signal would occur if it was present. We took a step size of 2% from the $0^{th}$ percentile to the $10^{th}$ percentile, and a step size of 10% from the $10^{th}$ percentile on to the $100^{th}$ percentile. The dashed horizontal line indicates the Null YFP Model's average performance with no ablations. The ML model's performance under ablation approaches the Null YFP Model's performance with no ablation at an intensity percentile of 30%. Error bars span one standard deviation in each direction above and below the average. (b) Example images are shown for both original YFP-tau images (leftmost column), YFP-tau images ablated at 30% (second column), original DAPI images (third column), and DAPI images ablated at 30% (rightmost column). We can see noticeable visual abnormalities at a high ablation of 30%. Images were auto-enhanced and colored with ImageJ and Matplotlib for visualization purposes only.

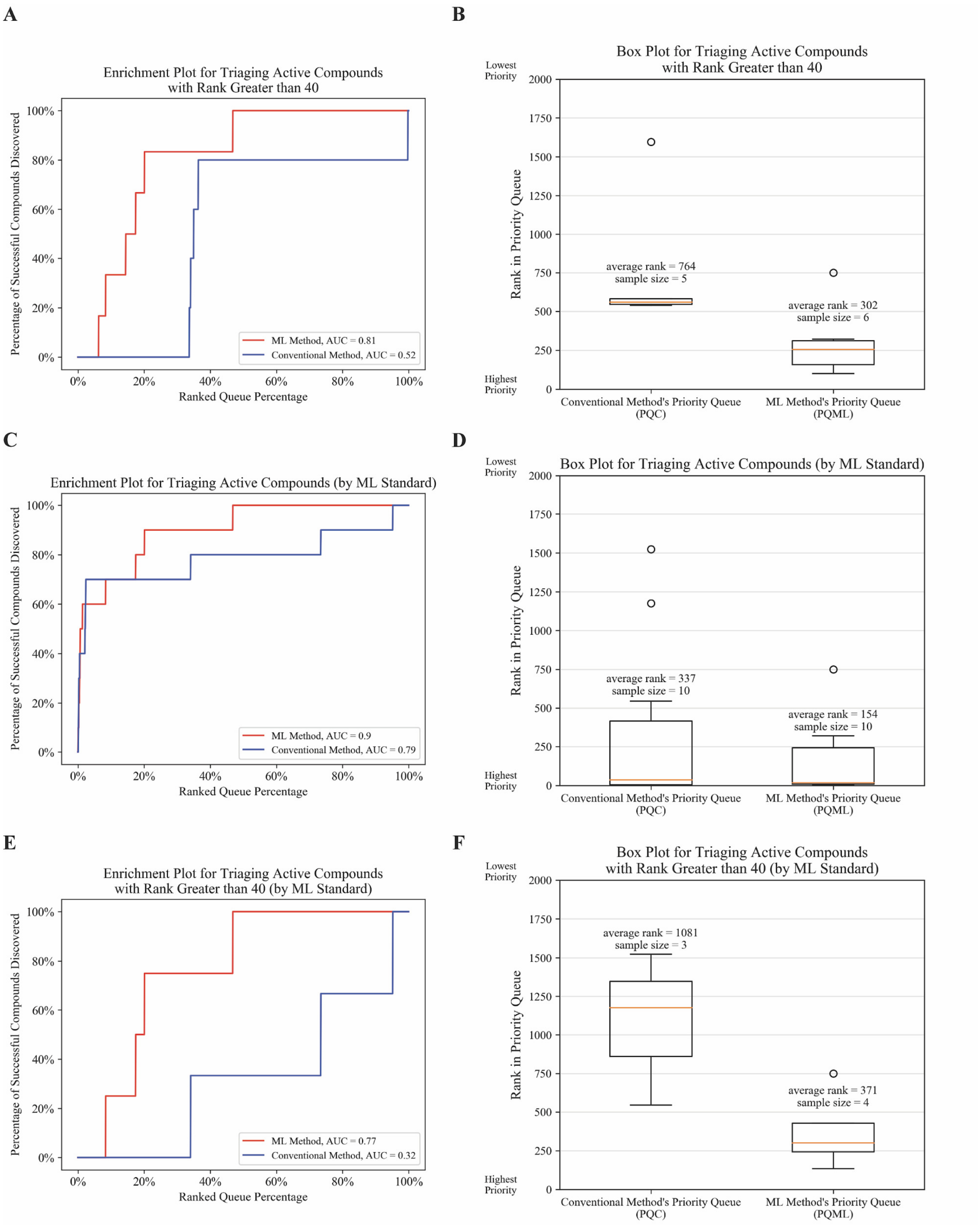


**Supplemental Figure 8: Additional triaging analysis of missed but active compounds, and also enrichment by a different ML Standard.** (a) An enrichment plot for active compounds that had rank greater than 40. (b) A box plot is depicted for active compounds residing outside of a queue's top 40 list. The ML achieved an average rank of 302, which was more than twice as low as the average rank of 764 for missed but active compounds in PQC. (c) - (f) correspond to enrichment analyses when assessing a compound's dose-response activity from the ML-derived AT8-pTau images instead of the YFP-tau images (denoted here as the "ML Standard"). Other than this difference in how images are scored for aggregation, the ML Standard is identical to the standard presented in the main text. (c) Enrichment curve for all active compounds by the ML Standard. By this standard, there were 10 active compounds. (d) Box plot for active compounds by the ML Standard. (e) Enrichment curve by the ML Standard for active compounds residing outside of the top 40 list. (f) Box plot by the ML Standard for active compounds residing outside of the top 40 list.

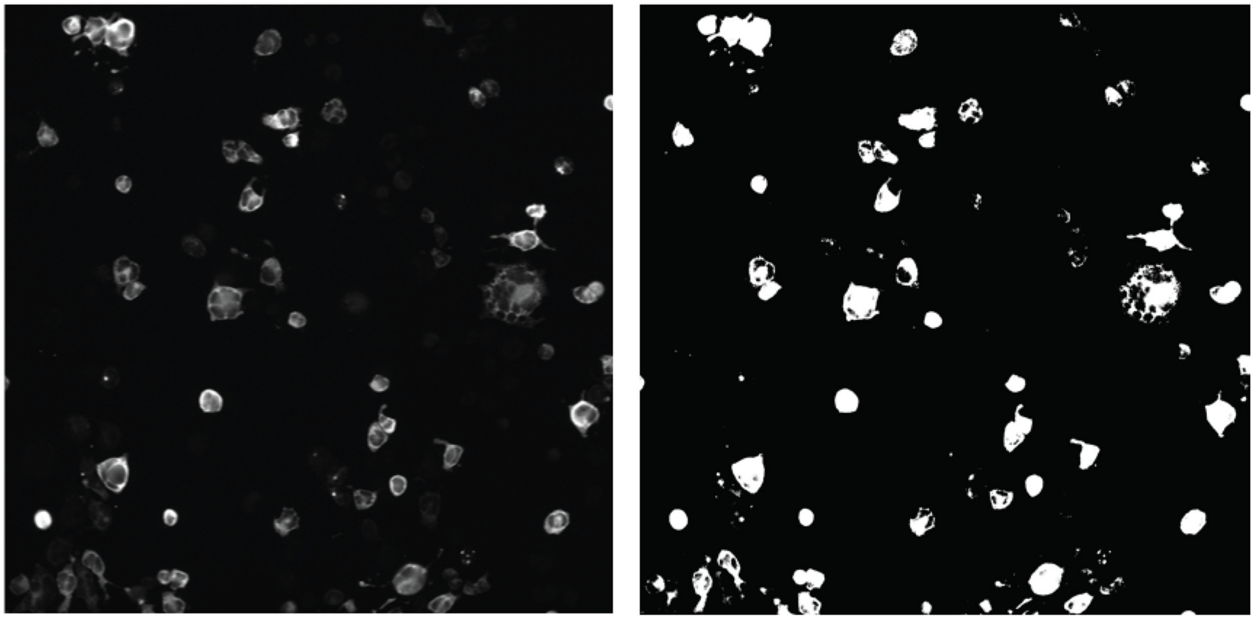

**Supplemental Figure 9: Binary thresholding of the label image in constructing the AUC.** The image on the left is the actual AT8-pTau label image (enhanced with ImageJ’s auto-enhance function for visualization purposes only). The image on the right is the result of normalizing the raw AT8-pTau image, and then thresholding with a pixel value = 1.0, such that any pixels greater than or equal to 1.0 are shown in white as positive signal, and any pixel with intensity less than 1.0 is shown in black as negative signal.
